# Off-targets of BRAF inhibitors disrupt endothelial signaling and differentially affect vascular barrier function

**DOI:** 10.1101/2023.08.24.554606

**Authors:** Sophie Bromberger, Yuliia Zadorozhna, Julia Maria Ressler, Silvio Holzner, Arkadiusz Nawrocki, Nina Zila, Alexander Springer, Martin Røssel Larsen, Klaudia Schossleitner

## Abstract

Targeted therapies against mutant BRAF are effectively used in combination with MEK inhibitors (MEKi) to treat advanced melanoma. However, treatment success is affected by resistance and adverse events (AEs). Approved BRAF inhibitors (BRAFi) show high levels of target promiscuity, which can contribute to these effects. Blood vessels are in direct contact with high plasma concentrations of BRAFi, but effects of the inhibitors in this cell type are unknown. Hence, we aimed to characterize responses to approved BRAFi for melanoma in the vascular endothelium. We showed that all clinically approved BRAFi induced a paradoxical activation of endothelial MAPK signaling. Moreover, phosphoproteomics revealed distinct sets of off-targets per inhibitor. Endothelial barrier function and junction integrity were impaired upon treatment with Vemurafenib and the next-generation dimerization inhibitor PLX8394, but not with Dabrafenib or Encorafenib. Together, these findings provide insights on the surprisingly distinct side effects of BRAFi on endothelial signaling and functionality. Better understanding of off-target effects could help to identify molecular mechanisms behind AEs and guide the continued development of therapies for BRAF-mutant melanoma.

## Introduction

Melanoma is a highly aggressive form of skin cancer and is associated with a high mortality rate (Schadendorf *et al*, 2018). Approximately 50% of melanomas harbor mutations in the *BRAF* gene, of which the vast majority encodes the BRAF-V600E oncoprotein (Davies *et al*, 2002; Ascierto *et al*, 2012). This mutation induces the constitutive activation of BRAF and downstream MAPK signaling and subsequently promotes excessive proliferation and survival of tumor cells. Targeted therapies, including inhibitors of mutant BRAF or its downstream effector MEK, are used to suppress this pathway in patients, with a combined approach yielding the best outcomes (Ascierto *et al*, 2016; Flaherty *et al*, 2012). Currently, three BRAF inhibitors (BRAFi) are clinically approved for the treatment of BRAF-V600E and BRAF-V600K mutant melanoma, namely Vemurafenib, Dabrafenib and Encorafenib, commonly administered together with the MEK inhibitors (MEKi) Cobimentinib, Trametinib, or Binimetinib, respectively (Larkin *et al*, 2014; Long *et al*, 2015; Dummer *et al*, 2018). While these targeted therapies have greatly improved the prognosis of patients with advanced BRAF-mutant melanoma, they also have two major limitations: On the one hand, acquired resistance to BRAF inhibition typically develops after a median of 9 - 12 months (Larkin *et al*, 2014; Long *et al*, 2015; Dummer *et al*, 2018). On the other hand, patients often experience adverse events (AEs), which lead to a discontinuation rate of up to 15.7% and to dose modifications in about 50% of patients (Heinzerling *et al*, 2019).

Numerous molecular processes potentially causing a resistance to BRAF inhibition have been studied and reviewed, among them MAPK-dependent and -independent mechanisms (Holderfield *et al*, 2014; Luebker & Koepsell, 2019). Yet the underlying molecular mechanisms for AEs remain largely unknown. It is often proposed, that resistance mechanisms as well as AEs can arise from a phenomenon called paradoxical ERK activation, which describes an activation of downstream MAPK signaling upon BRAF inhibition (Poulikakos *et al*, 2010; Adelmann *et al*, 2016). This phenomenon is caused by an alteration of RAS-dependent dimerization of BRAF (Lavoie *et al*, 2013). Newer drug development strategies include so-called “paradox breakers”, which are dimerization inhibitors designed to avoid paradoxical ERK activation (Brummer & McInnes, 2020). However, this inhibitor class still has to be clinically evaluated.

Furthermore, paradoxical activation of the MAPK pathway is not the only mechanism responsible for resistance and AEs. Therapeutic kinase inhibitors have repeatedly been investigated for their polypharmacology, meaning that they have binding capacities for a number of proteins aside from their designated target. For example, Vemurafenib has been shown to inhibit not only mutant BRAF, but also wildtype BRAF and CRAF in cell-free assays (Bollag *et al*, 2010). In higher concentrations it can inhibit a variety of other kinases, including LCK, YES1, SRC, or CSK. Dabrafenib has also been reported to act on wildtype BRAF and CRAF (Rheault *et al*, 2013). A comprehensive investigation on the target promiscuity of these inhibitors has been published in 2017, elucidating their binding capacities in protein lysates of cancer cells (Klaeger *et al*, 2017).

In recent years, it has been increasingly recognized that BRAFi have off-target effects on stromal cells of the tumor microenvironment (TME), such as fibroblasts, but also on immune cells, and that these off-targets could have a crucial impact on the treatment outcome (Corrales *et al*, 2021; Callahan *et al*, 2014; Loria *et al*, 2022). However, the vascular system has been severely underrepresented in this line of research, even though the vascular endothelium is in contact with high concentrations of BRAFi in the circulation. For example, patients receiving Vemurafenib experience plasma levels of up to 61.4 µg/ml (≙ 125.32 µM), which easily exceeds thresholds for interactions with multiple off-target kinases (Roche Registration Ltd., 2012; Bollag *et al*, 2010).

Impaired vascular function can be problematic, especially for patients with comorbidities (Lyon *et al*, 2020; Mincu *et al*, 2019). For example, endothelial dysfunction reduces the ability of blood vessels to dilate and can lead to increased peripheral resistance, a hallmark of hypertension (Vanhoutte *et al*, 2017; Ma *et al*, 2023). The endothelium helps to regulate the balance between pro- and anticoagulant mechanisms, and vascular damage can be a cause for disproportionate coagulation events, including thrombosis or hemorrhage (Neubauer & Zieger, 2022). Activation of adhesion receptors on endothelial cells can contribute to protumorigenic immune cell infiltrates and the formation of metastatic niches (Häuselmann *et al*, 2016; Wettschureck *et al*, 2019; Reymond *et al*, 2013). Increased permeability and the subsequent accumulation of excess fluid leads to higher interstitial pressure, which can limit treatment perfusion of the tumor and consequently can reduce therapeutic efficacy (Goel *et al*, 2011). No treatment against endothelial activation and vascular barrier disruption is available to date (Claesson-Welsh *et al*, 2021). Thus, in depth knowledge of the molecular signaling mechanisms in human endothelial cells is needed to inform future studies and therapeutic development.

In the present study, we aimed at elucidating the effects of BRAFi treatment on vascular endothelial signaling and functionality. We observed that paradoxical ERK activation occurs in endothelial cells. Simultaneously, numerous other signaling cascades were affected by BRAFi treatment, which we could show in a global mass spectrometry (MS)-based phosphoproteomics analysis. The comparison of several clinically used BRAFi revealed that endothelial off-targets were highly variable among treatments. Essential endothelial functions, most prominently the endothelial barrier, were also differentially affected by BRAFi treatment. Together, our data provide insights into the mechanisms of BRAFi-induced endothelial signaling disruption and dysfunction, which adds another piece to the puzzle of understanding the role of the TME in treatment outcomes and AEs in advanced melanoma.

## Results

### BRAFi induce paradoxical MAPK signaling in endothelial cells

Vemurafenib, Dabrafenib, Encorafenib and the next-generation dimerization inhibitor PLX8394 were designed to specifically target BRAF bearing the V600E mutation. However, to a certain extent, these inhibitors also act on wildtype BRAF and other off-targets in tumor cells in a concentration-dependent manner (Bollag *et al*, 2010; Rheault *et al*, 2013; Klaeger *et al*, 2017). In figure 1A, we could indeed show that increasing concentrations of Vemurafenib inhibited downstream ERK1/2 phosphorylation (T202/Y204) in primary BRAF-mutant melanoma cells and to a lesser degree in BRAF-wildtype melanoma cells after 1 h. In contrast, NRAS-mutant melanoma cells displayed a paradoxical activation of pERK after treatment with 1-10 µM Vemurafenib, which is in line with previous reports (Oh *et al*, 2016). Notably, when dermal microvascular endothelial cells (DMEC) were stimulated with the same concentrations of Vemurafenib, these cells also showed elevated pERK levels (Figure 1B and C). Likewise, this paradoxical activation could be seen after treatment of DMEC with low doses (1 µM) of Dabrafenib and Encorafenib (Figure 1D). Interestingly, the concentration at which we saw activation of ERK1/2 after Dabrafenib and Encorafenib was similar, but the peak of Vemurafenib-induced activation occurred at a higher concentration of 10 µM (Figure 1D). To investigate if the phosphorylation pattern in endothelial cells follows what is known for paradoxical activation of BRAF in melanoma cells, we added PLX8394, an inhibitor designed to avoid paradoxical activation, and indeed PLX8394-treated samples did not show elevated pERK levels in either cell type. All BRAFi were added at concentrations relevant for human use. While Vemurafenib has a maximum plasma concentration (C_max_) of up to 125.32 µM in clinical studies, standard dosing of Dabrafenib and Encorafenib led to plasma levels of 2.84 and 11 µM, respectively (Roche Registration Ltd., 2012; GlaxoSmithKline Trading Services Limited, 2013; Pierre Fabre Medicament, 2018). Although all BRAFi efficiently impeded ERK phosphorylation in BRAF-mutant and BRAF-wildtype melanoma cells, their effect on MAPK signaling in endothelial cells resembled the effect in melanoma cells with upstream NRAS mutations (Figure S1).

**Figure 1:**
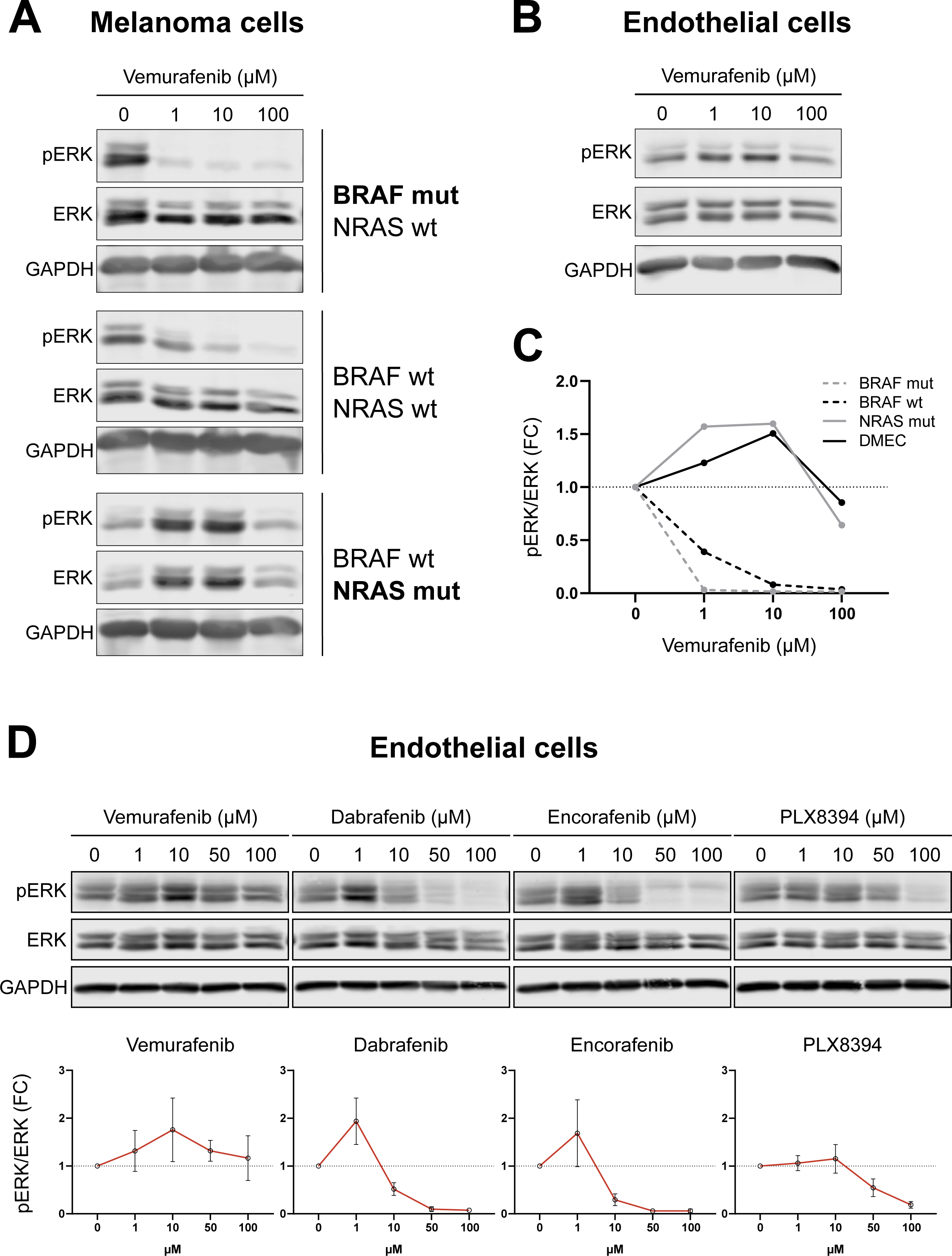
BRAFi induce paradoxical ERK activation in endothelial cells. A-B: Abundances of pERK (T202/Y204), total ERK and GAPDH in melanoma cells (A) and DMEC (B), treated with indicated concentrations of Vemurafenib for 1 h. C: Quantification of band intensities of shown blots (pERK/ERK ratio) displayed as fold change from the vehicle control. D: Western blots and respective quantifications of ERK phosphorylation in DMEC treated with indicated BRAFi concentrations for 1 h. Quantification of band intensities is displayed as fold change of pERK/ERK ratio from the vehicle control (mean ± SD, n = 3-4).

### Phosphoproteomics reveals BRAFi-induced disruption of endothelial signaling

To investigate if clinically relevant concentrations of BRAFi not only induce direct effects on MAPK signaling, but also induce off-target effects in endothelial cells, we utilized a mass spectrometry-based phosphoproteomics approach to determine altered phosphosites in phosphoproteins in DMEC after 1 h of BRAFi treatment. While BRAFi did not affect overall protein expression after 1 h (Figure S2), our analysis of phosphopeptides in figure 2A revealed distinct phosphorylation patterns for the tested inhibitors. The striking heterogeneity in phosphorylation among treatments became even more obvious when we found that most of the significantly altered phosphosites were unique for the individual inhibitors (Figure 2B). In more detail, only two phosphosites were commonly inhibited by all treatments, namely Desmoplakin (DSP, S2209) and Band 4.1-like protein 2 (EPB41L2, S87). A decrease in phosphorylation of Cortactin (CTTN, S261), Paxillin (PXN, S270), RHO GTPase-activating protein 29 (ARHGAP29, S356) and Liprin-beta-1 (PPFIBP1, S908) was observed in all treatments except 10 µM of Vemurafenib. We observed that Dabrafenib (10 µM) treatment led to the highest number of altered phosphosites, namely 107. Interestingly, the same concentration of Vemurafenib induced only minimal changes, whereas the higher concentration had stronger effects with 4 vs. 95 significantly altered phosphosites (Figure 2C). Reactome pathway enrichment analysis of the differentially phosphorylated proteins revealed that each of the inhibitors induced changes in its individual set of pathways (Figure 2D), which again highlighted the different global effects of BRAFi. For example, Vemurafenib (100 µM) interfered with pathways involved in mTOR and RHO GTPase signaling, whereas the effects of Dabrafenib were associated with different aspects of the MAPK pathway.

**Figure 2:**
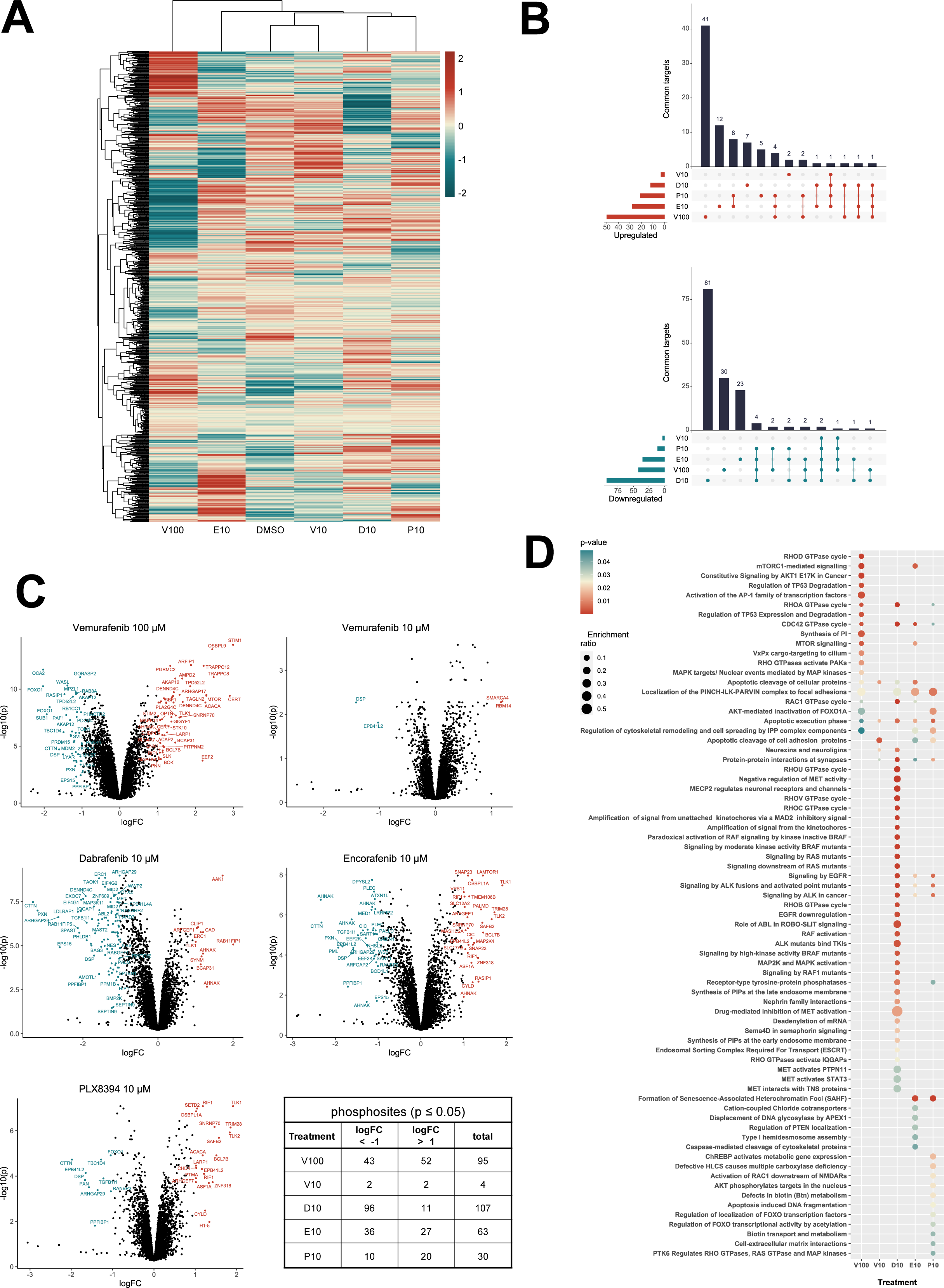
BRAFi disrupt the endothelial phosphoproteome. Mass spectrometry-based phosphoproteomics data of DMEC treated with vehicle control (DMSO), 10 µM of Vemurafenib (V10), Dabrafenib (D10), Encorafenib (E10), PLX8394 (P10), or 100 µM Vemurafenib (V100) for 1 h. A: Z-scored phosphosite abundance per condition (n = 3 donors). B: Overlaps of significantly up-(red) or downregulated (blue) phosphoproteins among treatments relative to the DMSO control (Limma, logFC ± 1, p ≤ 0.05). C: Phosphoprotein abundance of treatments compared to the DMSO control. D: Reactome pathway enrichment analysis of proteins with a significantly altered phosphorylation status, listed according to p-value and enrichment ratio calculated as entities found / total number of entities in the pathway.

We then integrated our experimental phosphopeptide abundance data with known kinase-substrate interactions to predict kinase activity via the previously published KinSwingR package (Engholm-Keller *et al*, 2019). Based on the phosphosites present in our dataset, we identified potential substrates for 156 kinases and computed their activity scores for each inhibitor treatment compared to the control. Hierarchical clustering of the 50 most differentially active kinases among inhibitor treatments emphasized the sometimes-contrasting effects of the used BRAFi on the vasculature (Figure 3A). STRING-based physical interaction networks of those kinases revealed that several CDKs and MAPKs, as well as GSK3-α/β were inhibited with Dabrafenib and the high dose of Vemurafenib, whereas Src-family kinases, AKT1 and protein kinases A and C were particularly activated with Dabrafenib (Figure 3B). The lower dose of Vemurafenib only differentially regulated four phosphoproteins and thus had only mild effects on kinase activation. Klaeger *et al* published an extensive study using cancer cell lysates to investigate the target promiscuity of 243 clinical kinase inhibitors, including Vemurafenib, Dabrafenib and Encorafenib (Klaeger *et al*, 2017). They employed a competitive affinity assay with immobilized broad-spectrum inhibitors (kinobeads) combined with mass spectrometry-based protein quantification, to assess which proteins would be bound by individual kinase inhibitors in lysates from leukemia, neuroblastoma and adenocarcinoma cell lines. We compared their datasets with the kinase activity predictions in our data from endothelial cells to deduce which off-targets could be directly bound by BRAFi and which could be downstream effectors (Table 1). Klaeger *et al* identified 10 proteins that were directly bound by Vemurafenib (up to 30 µM), three of which also occurred in our kinase dataset (BRAF, PTK6, TGFBR2). Notably, BRAF was paradoxically activated by 10 µM but inhibited by 100 µM Vemurafenib. We found an overlap of 24 kinases that were directly bound by Dabrafenib and were also present in our prediction dataset (total of 56 proteins in the Klaeger dataset). Especially a group of CDKs were strongly inhibited by Dabrafenib and also shown to be physically bound in low µM concentrations. Other kinases that were present in both datasets include ABL1/2, HCK, JAK2 and YES1. Interestingly, some kinases that were identified in the Klaeger dataset (CAMK4, FYN and LCK) were activated by Dabrafenib treatment. Encorafenib bound 28 proteins in the Klaeger dataset, 10 of which we also identified. In this case, especially GSK3-α/β as well as MAPK8/9 and MAPKAPK2 were inhibited in Encorafenib treated samples by direct interaction.

**Figure 3:**
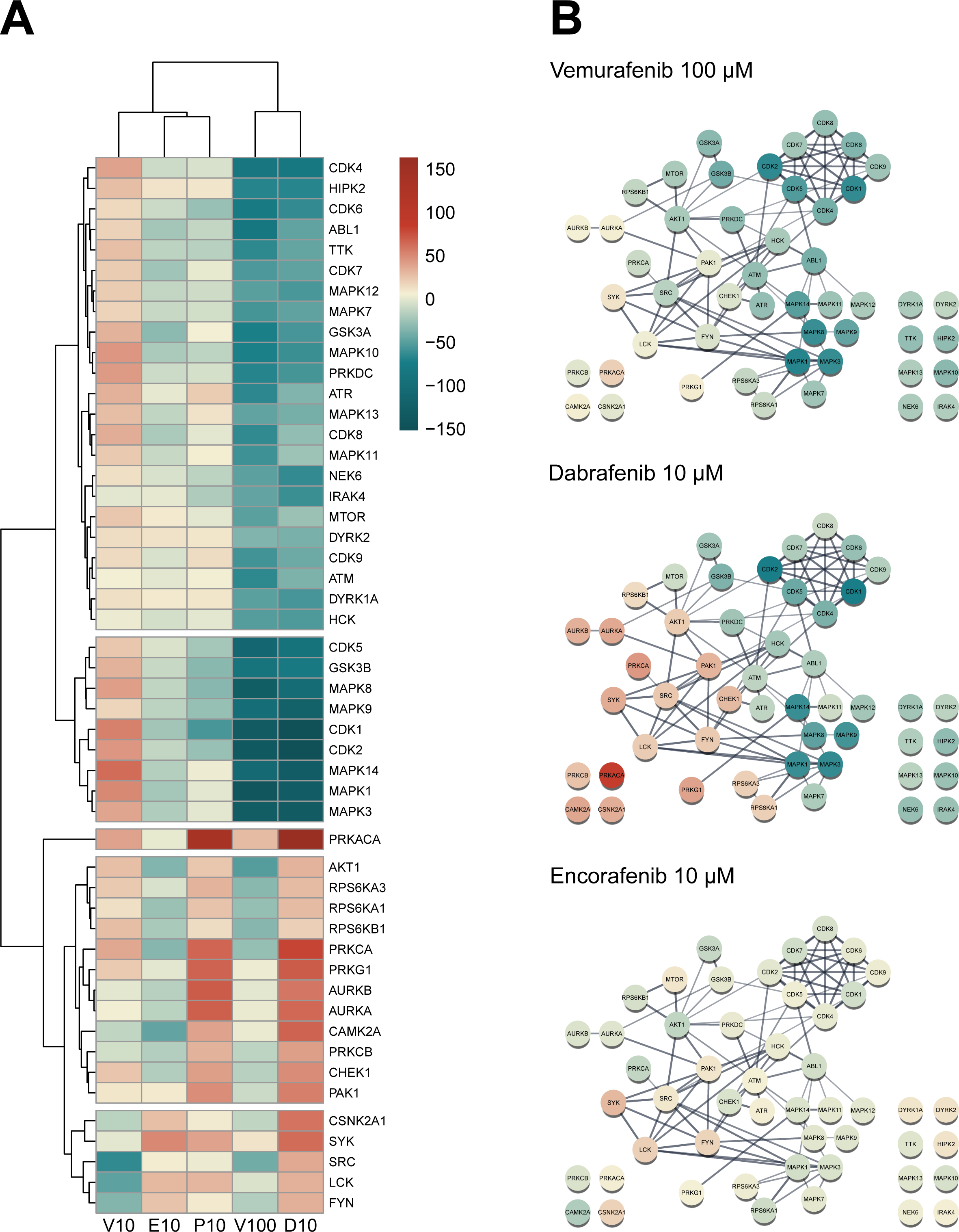
BRAFi differentially affect endothelial kinase signaling. A: Predicted kinase activity scores were computed from the phosphoproteomics dataset with KinSwingR. Weighted score for predicted activity of the 50 most differentially regulated kinases across all treatments compared to vehicle control. Scale ≙ Swing score. B: STRING physical subnetwork visualization of the same 50 kinases, Scale ≙ Swing score.

**Table 1:**
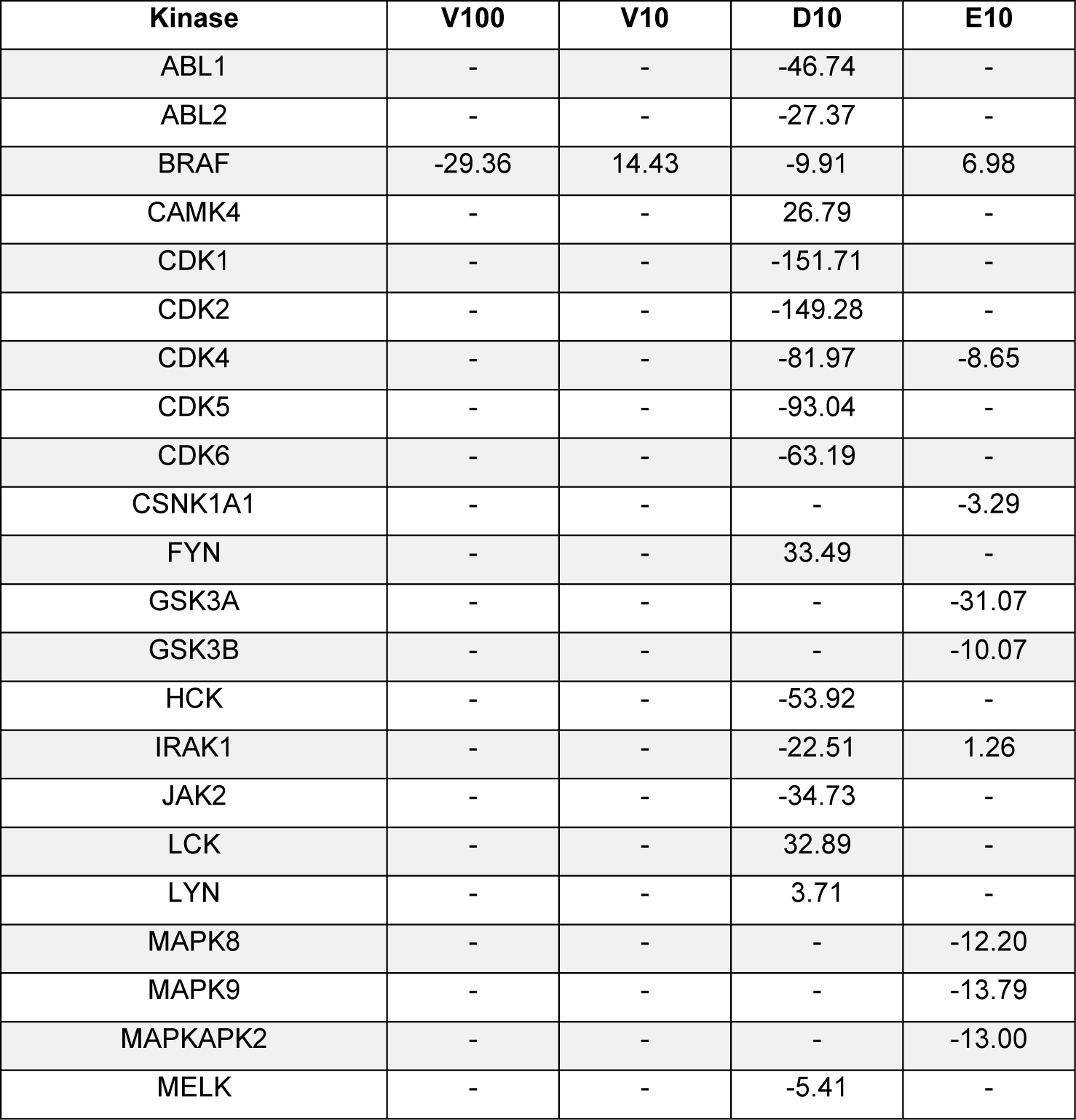
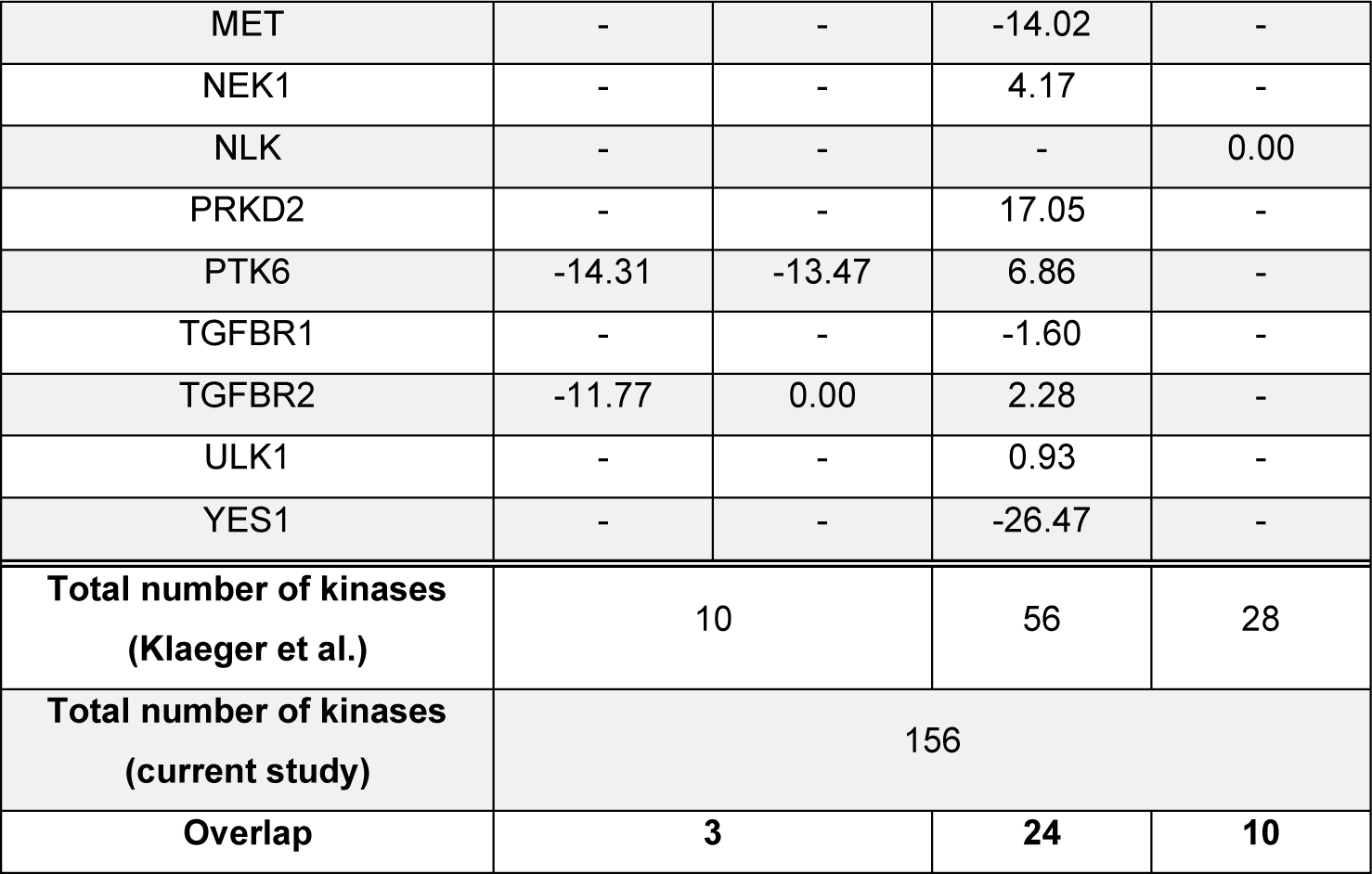
Comparison of KinSwing activity predictions with previously published data on direct target promiscuity of BRAFi (Klaeger *et al*, 2017). Predicted activity scores scaled according to Figure 3A are shown for kinases that were identified in both datasets. Empty cells indicate that the respective kinase was not directly bound in the Klaeger dataset. The last three rows indicate how many kinases were found per inhibitor in each dataset and how many were overlapping.

Our phosphoproteomics and kinase prediction data clearly highlight different off-targets among clinically used BRAFi in endothelial cells, even though these molecules were all designed to target mutant BRAF in melanoma cells.

### BRAFi differentially affect endothelial barrier function

Next, we investigated if the diverse effects on signaling pathways have functional consequences in endothelial cells. Endothelial morphology was not visibly altered by BRAFi (Figure S3A). Furthermore, they did not induce surface expression of activation markers such as ICAM-1 and E-Selectin (Figure S3B-C). However, electrical cell-substrate impedance sensing (ECIS) measurements of DMEC monolayers revealed a substantial dose-dependent disruption of electrical barrier resistance (measured at 250 Hz) by Vemurafenib (Figure 4A). Dabrafenib and Encorafenib did not show the same effect, even at high concentrations, whereas PLX8394 treatment induced a drop in barrier resistance. Concurrently, high doses of Vemurafenib and PLX8394 increased endothelial permeability of high (70 kDa) and low (376 Da) molecular weight tracers in a transwell assay after 1 h and 6 h when compared to the vehicle control (Figure 4B). Dabrafenib and Encorafenib had no effect on tracer permeability in this assay. In addition, we observed that high doses of Vemurafenib induced visible disruptions of endothelial cell-cell junctions (Figure 5A). Tight and adherens junctions appeared even and smooth in vehicle control-treated DMEC monolayers, as visualized by immunofluorescence of Claudin-5 and Vascular Endothelial (VE-)Cadherin. In contrast, junctions were interrupted and disorganized upon treatment with 100 µM of Vemurafenib. Lower doses (10 µM) of Vemurafenib or Dabrafenib did not alter the junction architecture to the same extent (Figure 5A). Similarly, 10 µM of Encorafenib and PLX8394 did not disturb junctions, but 100 µM of PLX8394 induced disruptions (Figure S4).

**Figure 4:**
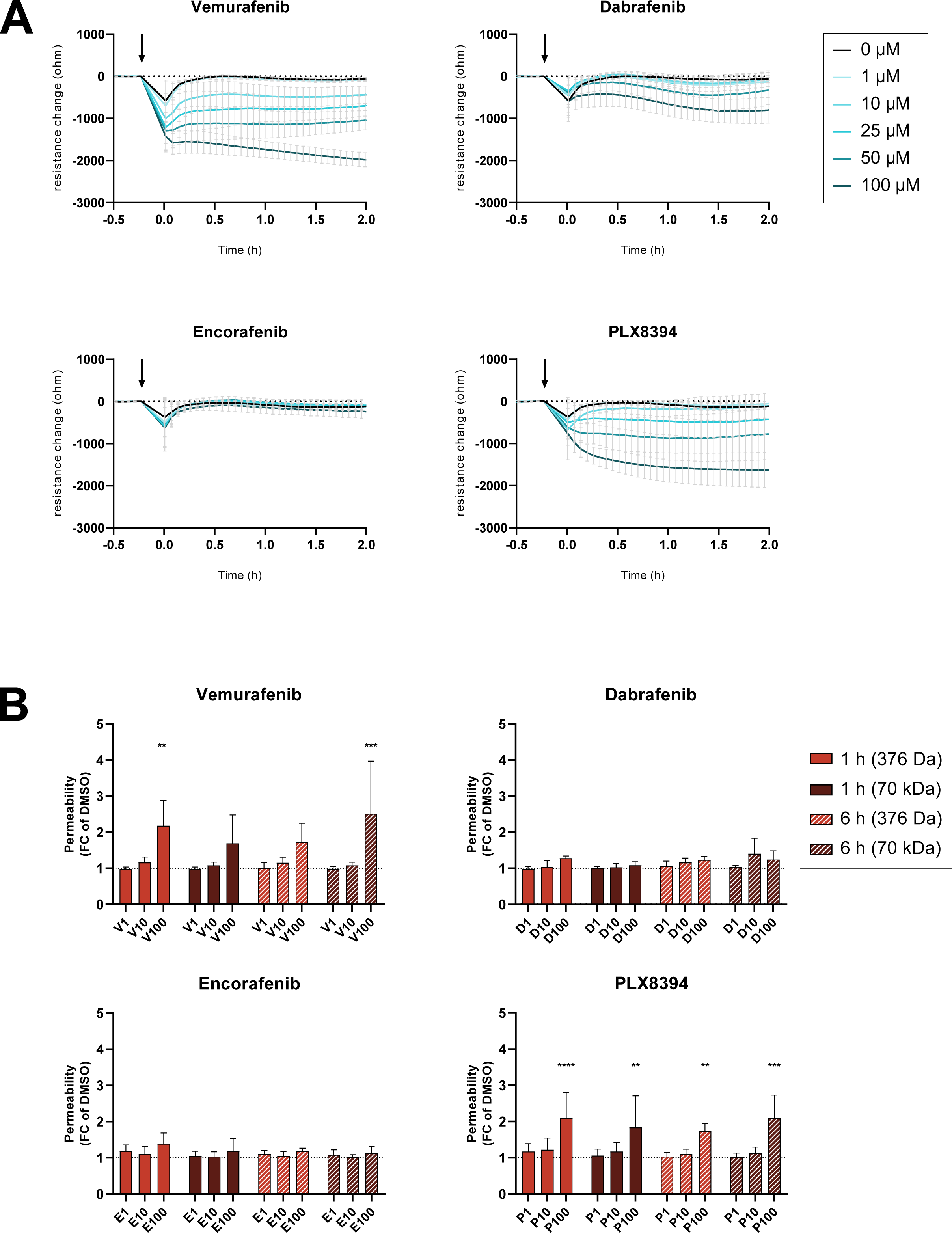
BRAFi differentially affect endothelial barrier function. A: ECIS real-time measurements of electrical barrier resistance in a DMEC monolayer upon BRAFi treatment, displayed as resistance change (ohm) from the time of inhibitor addition (mean ± SD, n = 5-10 biological replicates in four separate experiments). B: Permeability of fluorescently labelled tracers Na-Fluorescein (375 Da) and TRITC-dextrane (70 kDa) after 1 and 6 h of BRAFi treatment (n = 3-4 experiments with three biological replicates each). Results are depicted as mean ± SD. Significance was tested using two-way ANOVA and Dunnett’s test for multiple comparisons.

**Figure 5:**
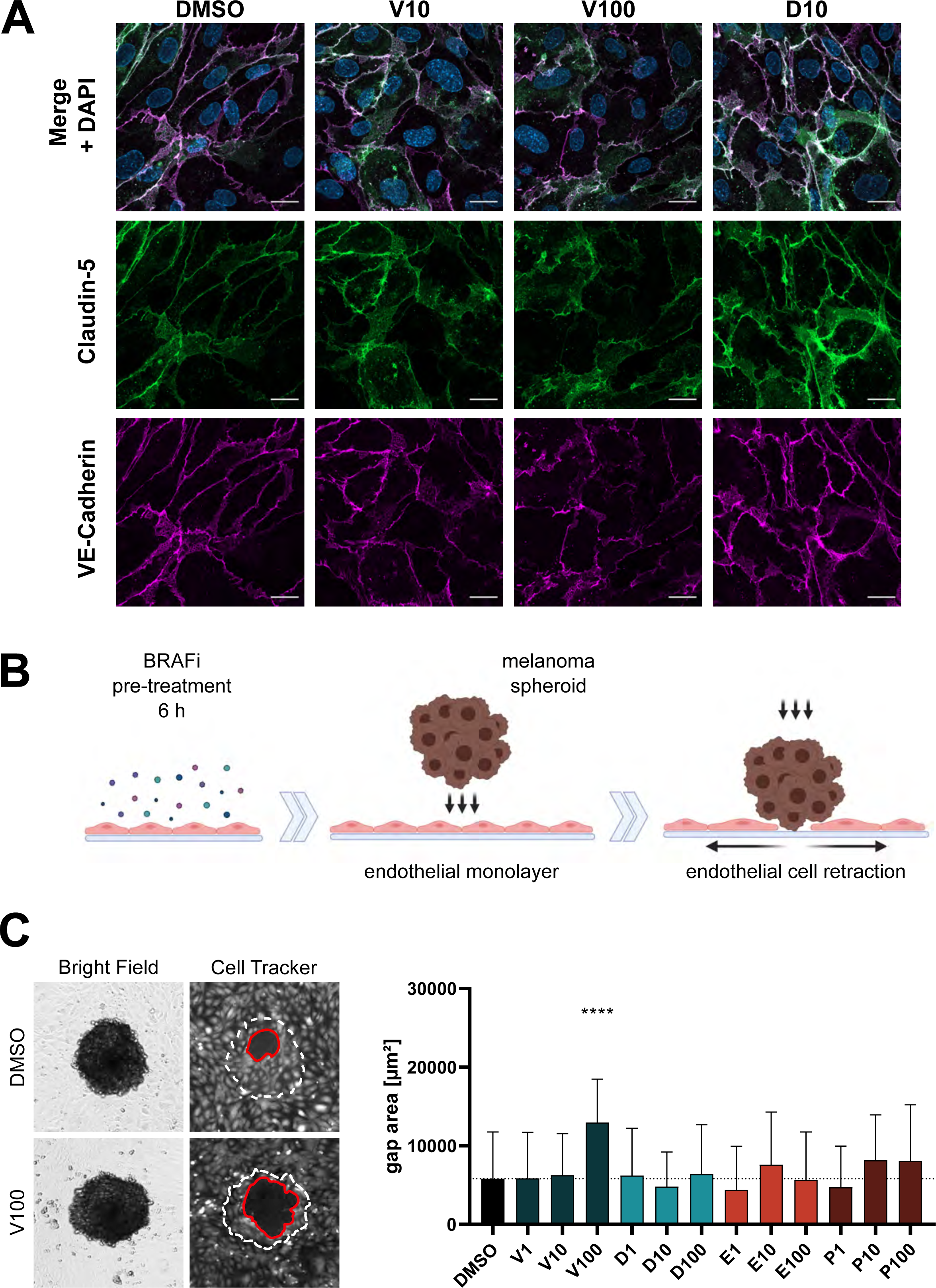
BRAFi differentially affect endothelial junctions and resistance to tumor cell invasion. A: Immunofluorescence of tight and adherens junctions in confluent DMEC, treated with DMSO, Vemurafenib, or Dabrafenib for 1 h. Green = Claudin-5, red = VE-Cadherin. Scale bars = 20 µm. B: Fluorescently labelled DMEC were treated with BRAFi for 6 h, prior to incubation with melanoma spheroids for 6 h. C: The area of gaps (red line) in the endothelial monolayer beneath spheroids (white dotted line) is depicted as mean ± SD (n(treatment) = 37-60 spheroids, n(control) = 176 spheroids, from at least three independent experiments).

A weak endothelial barrier can have detrimental consequences, not only because of the leakage of fluid and small molecules, but also during metastasis formation. Based on a previously published *in vitro* model of tumor cell invasion (Holzner *et al*, 2016), we measured the size of melanoma spheroid-induced gaps in BRAFi-treated DMEC monolayers (Figure 5B). Pre-treatment with high doses of Vemurafenib weakened the endothelial barrier against invading tumor cells, resulting in a significantly larger gap area compared to vehicle control treatment (Figure 5C). None of the other inhibitors affected the spheroid-induced gap area.

### BRAFi affect vascular junctions in patients

We further aimed to investigate the effects of clinically used BRAFi on the vasculature in skin biopsies from advanced melanoma patients who had been treated with BRAFi. We obtained archived skin biopsies before and during therapy from one patient who had received Vemurafenib monotherapy, one with Vemurafenib + Cobimetinib, and three patients who had been treated with Dabrafenib + Trametinib and from their matched control. Tissue sections were then subjected to immunofluorescence staining for vascular markers. Our image analysis showed that within VE-Cadherin-positive vessels, the signal of the tight junction protein Claudin-5 was decreased upon Vemurafenib monotherapy (72.44% during treatment vs. before), whereas the combinations of Vemurafenib + Cobimetinib and Dabrafenib + Trametinib did not have strong effects (Table 2, Figure 6). In Podoplanin-positive lymphatic vessels, we found a decrease in both VE-Cadherin and Claudin-5 signal upon Vemurafenib monotherapy (62.47% and 40.96% during treatment vs. before, respectively). This effect was not observed in patients who received either of the combination treatments. Of note, patient 5, who was matched with a control sample from a different patient, generally displayed higher fluorescence intensity values during treatment for all markers and quantification masks.

**Figure 6:**
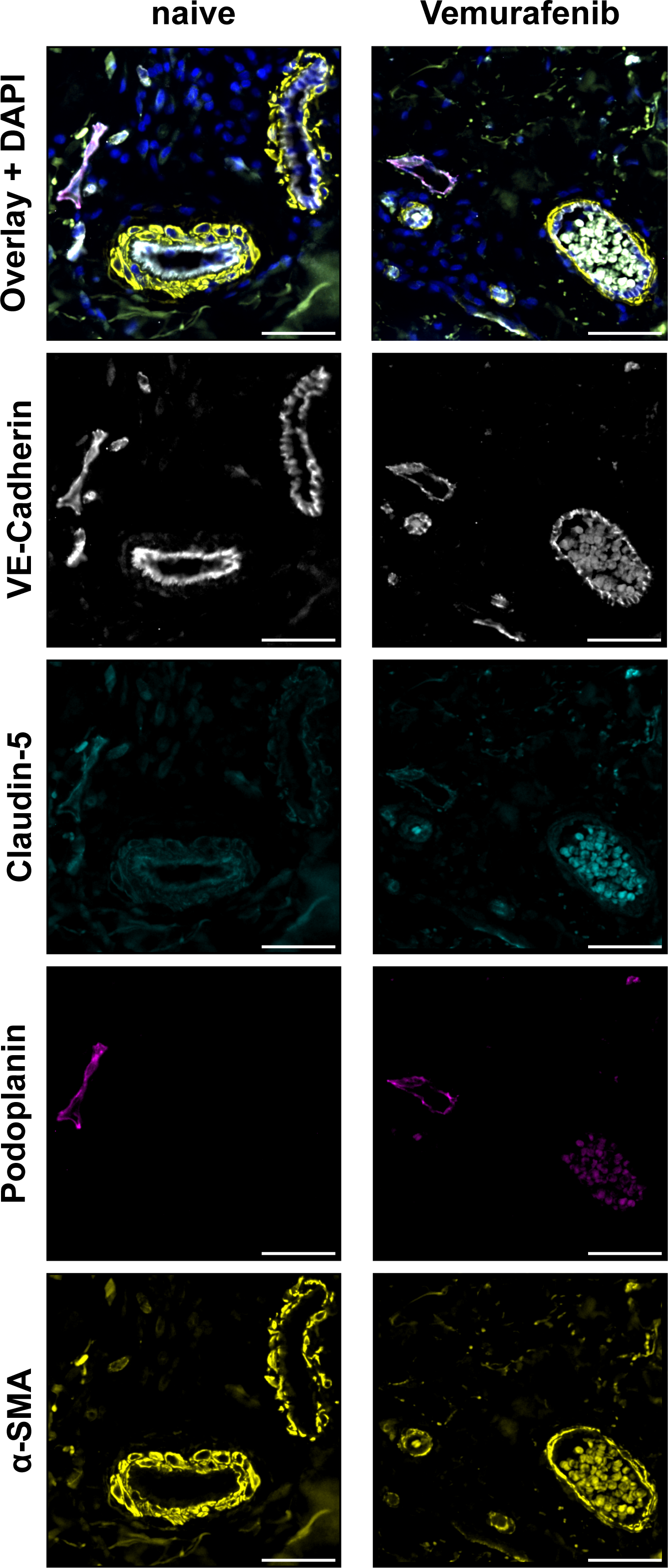
Effect of Vemurafenib on patient vessels. Immunofluorescence images of vascular markers in skin biopsy sections of one melanoma patient before (naïve) and during Vemurafenib monotherapy. Markers are VE-Cadherin (white), Claudin-5 (cyan), Podoplanin (magenta) and α-smooth muscle actin (yellow). Scale bars = 50 µm.

**Table 2:** Vascular junction markers in skin biopsies before and during treatment with indicated inhibitors. Fluorescence signal intensities of vascular markers during treatment are displayed as % of the intensity values before treatment within the same patient, or a matched control. Values were quantified within areas that were positive for VE-Cadherin or Podoplanin, to specifically measure intensities within all vessels or lymphatic vessels, respectively. Overall intensity refers to the mean fluorescence within the entire region of interest, including background and the stromal compartment.

| Patient # | 1 | 2 | 3 | 4 | 5 |
| --- | --- | --- | --- | --- | --- |
| Treatment | Vemurafenib monotherapy | Vemurafenib + Cobimetinib | Dabrafenib + Trametinib | Dabrafenib + Trametinib | Dabrafenib + Trametinib |
| VE-Cadherin<sup>+</sup> vessels |  |  |  |  |  |
| Claudin-5 (%) | 72.44 | 93.28 | 91.73 | 92.08 | 180.41 |
| Podoplanin<sup>+</sup> vessels |  |  |  |  |  |
| VE-Cadherin (%) | 62.47 | 133.33 | 130.86 | 118.09 | 168.88 |
| Claudin-5 (%) | 40.96 | 121.72 | 106.21 | 98.22 | 221.40 |
| Overall intensity across all pixels |  |  |  |  |  |
| VE-Cadherin (%) | 128.78 | 93.96 | 98.38 | 150.72 | 103.94 |
| Claudin-5 (%) | 134.05 | 98.15 | 86.05 | 178.97 | 110.94 |
| Podoplanin (%) | 120.76 | 96.43 | 95.14 | 120.62 | 122.75 |
| Total area for analysis |  |  |  |  |  |
| before treatment (μm<sup>2</sup>) | 58 019 379 | 62 369 079 | 25 270 836 | 12 909 993 | 12 445 770 |
| during treatment (μm<sup>2</sup>) | 9 740 901 | 30 001 823 | 6 542 780 | 4 372 562 | 11 464 887 |

## Discussion

Targeted therapies aimed at mutant BRAF and downstream MAPK signaling components are effective treatments against BRAF-V600E/K-positive melanoma. Currently, three different inhibitors against advanced BRAF-mutant melanoma are clinically approved for therapy: Vemurafenib, Dabrafenib and Encorafenib. They have different pharmacodynamic profiles, that are connected to their inhibitory potency towards mutant BRAF and off-target effects. A detailed review from 2019 discusses the differences among clinically used BRAF and MEK inhibitors, regarding their pharmacodynamics and particularly their adverse event profiles (Heinzerling *et al*, 2019). Vascular endothelial cells come in direct contact with high inhibitor concentrations and could influence treatment outcomes, however, studies that investigate the effects of BRAFi on the vascular endothelium are lacking. Therefore, our findings of differential effects of clinically used BRAFi on endothelial cells inform the field about potential pathways that could elicit side effects or provide a protumorigenic microenvironment.

### Paradoxical MAPK activation

We observed that all of the inhibitors approved for clinical use increased ERK phosphorylation in DMEC in a concentration-dependent manner, which corresponds to clinical dosing: The C_max_ of Vemurafenib (61.4 µg/ml ≙ 125.32 µM) is approximately 40 times higher than that of Dabrafenib, and 10 times higher than Encorafenib (Roche Registration Ltd., 2012; GlaxoSmithKline Trading Services Limited, 2013; Pierre Fabre Medicament, 2018). For endothelial cells the concentration of BRAFi measured in the patient circulation is critical. The current treatment regimen for BRAFi is not adjusted according to the weight or biological sex of the patient (Garbe *et al*, 2022). This has been reported as a potential factor for increased AEs and dose modifications, especially in women or patients with low body weight (Hopkins *et al*, 2020). However, these groups also experienced a benefit from a higher exposure to BRAFi and have shown higher overall survival (Vellano *et al*, 2022). In our study, we could show that clinically relevant BRAFi concentrations represent a sensitive balance between paradoxical activation and inhibition of the MAPK pathway in endothelial cells: In cell culture, concentrations of 10 µM Vemurafenib were necessary to induce paradoxical ERK activation. In comparison, Dabrafenib and Encorafenib induced ERK phosphorylation already at a lower dose of 1 µM. This paradoxical MAPK activation did not occur in DMEC treated with the so-called “paradox breaker” PLX8394, a next-generation BRAFi designed to specifically interfere with the dimerization dynamics of mutant BRAF (Basile *et al*, 2014). The phenomenon of paradoxical MAPK activation has been extensively investigated in BRAF wildtype cancer cells, especially in the presence of upstream NRAS mutations (Hatzivassiliou *et al*, 2010; Oh *et al*, 2016). In recent years, there have been increased efforts to elucidate the response to BRAFi not only in tumor cells, but also in other cells in or outside of the tumor microenvironment (TME). For example, BRAFi-induced paradoxical ERK activation has been shown in fibroblasts, keratinocytes and immune cells (Corrales *et al*, 2021; Escuin-Ordinas *et al*, 2016; Callahan *et al*, 2014). This has cell-type specific functional consequences and could play a crucial part in the outcome of BRAFi treatment. Previous publications suggested, that the effects of BRAFi on the TME could be contributing to melanoma clearance by improving T cell infiltration (Knight *et al*, 2013; Wilmott *et al*, 2012). However, a more recent study showed that paradoxical MAPK signaling in macrophages of the TME could also have tumor-protective effects by promoting resistance mechanisms (Wang *et al*, 2015).

### Off-targets in endothelial cells

Vemurafenib is known to target not only mutant BRAF, but also CRAF, SRMS and ACK1 with a similar IC_50_ (18 – 48 nM) in cell-free assays (Bollag *et al*, 2010). At concentrations in the low µM range it inhibits numerous other kinases. The plasma levels of Vemurafenib are by far exceeding thresholds for interfering with a broad range of kinases. This suggests that apart from MAPK other signaling pathways would also be affected by BRAFi treatment. To gain deeper insights into kinase signaling dynamics of BRAFi treatment, we performed mass spectrometry-based proteomics and phosphoproteomics of DMEC treated with the respective inhibitors. We observed no changes in protein abundance, but all used BRAFi had considerable effects on phosphorylation after 1 h of treatment. To our surprise, each BRAFi affected a specific subset of phosphoproteins in and outside of the MAPK pathway. Vemurafenib in the lower concentration (10 µM) caused only minor alterations in the phosphoproteome of endothelial cells, but the higher dose of 100 µM had a similarly strong effect as 10 µM of the other BRAFi, which also underlines the pharmacodynamic differences among these inhibitors. The different effects of BRAFi suggest that phosphosites are altered by off-target kinases outside of the MAPK pathway. This hints at a considerable amount of polypharmacology, or target promiscuity, which describes the capacity of an inhibitor to bind more than one target. Target promiscuity can be attributed to the fact that most kinase inhibitors attack the ATP-binding pocket of their target, which is structurally similar among kinases and other enzymes (Karoulia *et al*, 2017; Tong & Seeliger, 2015). This phenomenon can have detrimental but also beneficial aspects, especially in drug repurposing, but also in complex diseases such as cancer, where concomitant manipulation of oncogenic pathways could either impede or improve treatment efficacy (Kabir & Muth, 2022). For example, recent publications have identified mTOR signaling and the SEMA6A/RHOA/YAP axis as off-target mechanisms in BRAFi-associated tumor-protective effects of fibroblasts in the TME (Seip *et al*, 2016; Loria *et al*, 2022). The relevance of effects in the TME is evident and we are first to describe the molecular consequences of second and third-generation BRAFi treatment on the vascular endothelium.

To truly understand the promiscuous nature of therapeutic agents, a comprehensive analysis of on- and off-target effects is necessary, in which proteomics and PTMomics play a central role (Zecha *et al*, 2023). A study by Klaeger *et al* investigated the target promiscuity of clinical kinase inhibitors with a competitive affinity assay (kinobeads) paired with mass spectrometry to assess which proteins would be bound by individual kinase inhibitors in cancer cell lysates (Klaeger *et al*, 2017). Comparing their datasets with our kinase activity predictions in endothelial cells, we found notable parallels between physical binding and activity regulation for Vemurafenib, Dabrafenib and Encorafenib. Although physical binding affinity correlated with kinase inhibition in most cases, some of the kinases that were bound by Dabrafenib in the Klaeger dataset, including CAMK4, FYN and LCK showed a higher predicted activity in endothelial cells. Additionally, BRAF activity was increased after treatment with 10 µM, but decreased with 100 µM Vemurafenib, highlighting a dynamic dose response that could also apply to off-target kinases. A comparison of these two datasets highlights two aspects of BRAFi dynamics in human cells: Predicted kinase activity in an intact layer of live endothelial cells complements physical binding data in cancer cell lysates. It allowed us to identify inhibiting and activating off-targets that overlapped between the datasets. Together with the above cited published studies about off-targets of BRAFi, our findings provide additional support to the hypothesis that effects cannot be attributed to aberrant MAPK signaling alone, but also arise from target promiscuity.

The heterogenous alterations of protein targets among different BRAFi were also reflected in our analysis of enriched signaling pathways in BRAFi-treated endothelial cells. We observed that each BRAFi manipulated its individual set of pathways. For example, we observed enriched terms involving RHO GTPase signaling particularly in samples treated with 100 µM Vemurafenib. The importance of RHO GTPases in endothelial homeostasis, especially in angiogenesis and permeability, has been discovered many years ago (Van Nieuw Amerongen *et al*, 2000; Carbajal & Schaeffer, 1999; Wojciak-Stothard *et al*, 1998). It is known that RHOA regulates vascular permeability by interacting with the cytoskeleton at the site of endothelial junctions, which destabilizes cell-cell contacts (van Buul & Timmerman, 2016; Reinhard *et al*, 2017). Surprisingly, only Dabrafenib treatment was associated with enriched terms regarding RAS and RAF signaling. These findings highlight the distinct off-target effects of BRAFi in endothelial cells.

### Functional implications

There is not much literature regarding functional effects of BRAFi on endothelial cells, apart from one recent publication that investigated the effect of 28 clinically used kinase inhibitors on endothelial permeability (Dankwa *et al*, 2021). However, they did not include any BRAF-specific inhibitors, except the first-generation inhibitor Sorafenib, which induced a weakly barrier-disruptive phenotype. A plasma proteome analysis of melanoma patients reported increased plasma levels of KDR (VEGFR2) upon treatment with BRAFi (Babačić *et al*, 2021). Although it was not further discussed, this could hint towards increased VEGFR2 shedding and therefore dysregulated VEGF-A signaling, or endothelial dysfunction. Clinical studies have shown that combined inhibition of VEGF and BRAF delayed the onset of acquired resistance to Vemurafenib (Comunanza *et al*, 2017).

In this study, we observed that Vemurafenib and PLX8394 induced a dose-dependent breakdown of electrical barrier resistance as well as a hyperpermeability for low and high molecular weight fluorescent tracers. High doses also interrupted the integrity of endothelial tight and adherens junctions. Thus, our functional results confirmed what would be expected from our phosphoproteome analysis and the above cited literature, especially for altered RHO GTPase signaling. However, we did not observe a correlation between paradoxical ERK activation and functional response. ERK signaling was shown to play an important role in vascular integrity and endothelial knockout of *Erk2* in adult *Erk1*^-/-^ mice was associated with lethality due to vascular defects across all organ systems (Ricard *et al*, 2019). All clinically approved inhibitors induced a paradoxical ERK activation, but only Vemurafenib treatment caused severe endothelial barrier dysfunction. Additionally, the dimerization inhibitor PLX8394 did not induce paradoxical MAPK signaling but had similar effects on endothelial function as Vemurafenib. Despite the widespread assumption that paradoxical MAPK signaling is mainly responsible for unwanted side effects from data in stromal cells (Adelmann *et al*, 2016), we provide proof that this is not the case for vascular endothelial cells.

In addition, we investigated the barrier resistance of endothelial cells against tumor cell spheroids based on a previous model of tumor cell invasiveness (Holzner *et al*, 2016). In our experimental setup, only a high dose of Vemurafenib significantly weakened the endothelium against melanoma cell spheroids. Although this simplified model does not account for many factors involved in metastasis formation *in vivo*, it gives insights about Vemurafenib in facilitating the transmigration of tumor cells. Indeed, it has been previously shown that Vemurafenib treatment was associated with a higher metastatic burden in a drug-resistant melanoma mouse model (Obenauf *et al*, 2015).

It is important to note, that Vemurafenib is barely prescribed to patients nowadays, because Dabrafenib and Encorafenib, together with their respective MEKi, exhibit superior response rates and toxicity profiles, while showing a reduced occurrence of secondary neoplasms, when compared to Vemurafenib (Garbe *et al*, 2022; Heinzerling *et al*, 2019). Current clinical guidelines recommend targeted therapies as a second-line treatment after immune checkpoint inhibitor (ICI) therapy for advanced melanoma, although the optimal sequencing strategy for different patient groups is still investigated (Keilholz *et al*, 2020). Interestingly, the next-generation dimerization inhibitor PLX8394 showed very promising results in preclinical studies, but no clinical data are available to date. A Phase I/IIa trial (NCT02012231) was registered in late 2013 for evaluating the safety and preliminary efficacy of PLX8394, but never moved forward to Phase 2. Another Phase I/IIa trial (NCT02428712) was registered in 2015, but has not been completed yet. Notably, Dabrafenib and Encorafenib had no effect on the endothelial barrier function even at high concentrations, despite inducing paradoxical MAPK activation and affecting multiple off-target pathways.

### Clinical consequences

The AEs documented for clinically approved BRAFi are distinct from one another, especially when combined with their corresponding MEKi. A number of cutaneous and gastrointestinal AEs are common among all inhibitors, while other events occur more frequently with particular BRAFi or MEKi. For example, QT prolongation was observed in up to 7% of patients undergoing Vemurafenib monotherapy (Heinzerling *et al*, 2019; Flaherty *et al*, 2014), whereas other cardiovascular events such as pulmonary embolism, arterial hypertension and decreased left ventricular ejection-fraction have been linked to MEKi (Mincu *et al*, 2019; Abdel-Rahman *et al*, 2016). Vasculitis has been described sporadically as an AE in Vemurafenib-treated patients (Heinzerling *et al*, 2019).

We aimed to translate our findings from cell culture into a clinical context by investigating junctional markers of dermal vessels of pre- and on-treatment biopsies from cutaneous metastases of melanoma patients who had received BRAFi therapy. Our inclusion and exclusion criteria yielded five eligible patients who were treated at our clinic either with Vemurafenib monotherapy (one patient), Vemurafenib + Cobimetinib (one patient), or Dabrafenib + Trametinib (three patients). Immunofluorescence showed a decrease in endothelial junction markers (VE-Cadherin, Claudin-5) upon Vemurafenib monotherapy, whereas the combination treatments did not have the same effect. These results demonstrate vessel damage upon Vemurafenib therapy in one patient, which is coherent with our findings in cultured human endothelial cells. Thus, detrimental effects on endothelial cells could be a potential explanation for the higher AE rate in Vemurafenib compared to other BRAFi treated patients. However, further studies with larger sample sizes are needed to specifically characterize vascular-specific effects of new BRAFi and their consequences for AEs in patients.

In conclusion, we present evidence that inhibitors against mutant BRAF have considerable effects on the vascular endothelium. Although all clinically approved BRAFi induced paradoxical MAPK activation in endothelial cells, their off-target spectra are diverse. This is also reflected in their functional impact on the endothelium. Especially Vemurafenib substantially disrupted endothelial barrier function. Therefore, together with the off-target profiles acquired by phosphoproteomics, our results provide proof that BRAFi disrupt endothelial homeostasis. This could give insights into the mechanisms that are responsible for AEs. Future therapeutic developments and clinical studies should consider the target promiscuity of kinase inhibitors in the tumor microenvironment, including the vasculature. Better knowledge of the response to BRAFi in tumor cells and cells of the TME seems critical for future developments and could help to find even better treatment options for specific patient groups.

## Materials & Methods

### Antibodies and reagents

The BRAFi used in this study were Vemurafenib (S1267), Dabrafenib (S2807), Encorafenib (S7108) and PLX8394 (S7965), all purchased from Selleckchem (Houston, TX, USA). DMSO (D2650, Sigma-Aldrich, St. Louis, MO, USA) at a concentration of 0.1% was applied as a vehicle control. All antibodies used for this study are presented in Table 3.

**Table 3:** Primary and secondary antibodies used in the present study were purchased from the following suppliers: CST = Cell Signaling Technologies (Danvers, MA, USA), abcam (Cambridge, UK), LI-COR (Lincoln, NE, USA), Beckman Coulter (Brea, CA, USA), Invitrogen (Waltham, MA, USA). Antibody dilutions are indicated per method as WB for Western Blot and IF for immunofluorescence.

| Target | Supplier | Cat. Nr. | Dilution |
| --- | --- | --- | --- |
| p-ERK1/2 (T202/Y204) | CST | 9101 | 1:1,000 (WB) |
| ERK1/2 | CST | 4695 | 1:1,000 (WB) |
| GAPDH | abcam | ab181602 | 1:20,000 (WB) |
| Rabbit IgG (IRDye® 800CW-conjugated 2<sup>nd</sup> step) | LI-COR | 925-32213 | 1:15,000 (WB) |
| Rabbit IgG (IRDye® 680RD-conjugated 2<sup>nd</sup> step) | LI-COR | 925-68073 | 1:15,000 (WB) |
| VE-Cadherin | Beckman Coulter | IM1597 | 1:200 (IF cells) |
| VE-Cadherin | CST | 93467 | 1:200 (IF tissue) |
| Claudin-5 (AF488-conjugated) | Invitrogen | 352588 | 1:200 (IF cells) |
| Claudin-5 | Invitrogen | 352500 | 1:200 (IF tissue) |
| α-SMA (FITC-conjugated) | Sigma-Aldrich | F-3777 | 1:5,000 (IF tissue) |
| Podoplanin (AF647-conjugated) | Biolegend | 337008 | 1:500 (IF tissue) |
| Mouse IgG (AF546-conjugated 2<sup>nd</sup> step) | Life technologies | A-11030 | 1:500 (IF cells/tissue) |
| Rabbit IgG (AF594-conjugated 2<sup>nd</sup> step) | Life technologies | A-32754 | 1:500 (IF tissue) |

### Cell culture

Human DMEC were isolated from freshly discarded foreskin of pediatric patients undergoing circumcisions at the Department of Pediatrics of the Medical University of Vienna. The protocol was approved by the Ethics Committee of the Medical University of Vienna (1621/2020) in accordance with the Declaration of Helsinki. Written informed consent was obtained from the patients’ legal guardians. The collected tissues were cut into thin strips before incubation with Dispase (CLS354235, Merck, Darmstadt, Germany) for 20 min at 37°C. After gentle elimination of the epidermal layer, cells were dislodged with a cell scraper in Endothelial Cell Growth Medium MV (EGM-MV; C-22020, Promocell, Heidelberg, Germany) with the respective supplement mix, 15% fetal calf serum (FCS; 10500-064, Gibco, Karlsruhe, Germany), and 50 µg/ml Gentamicin (15710-049, Gibco). The cell suspension was centrifuged at 800 rpm for 10 min, pelleted cells were resuspended in fresh medium and seeded onto a 6-well plate. Until the first passaging, 100 µg/ml primocin (ant-pm-1, InvivoGen, San Diego, CA, USA) was added to the culture medium. Prior to the first passaging, cells were sorted with Dynabeads for CD31 (11155D, Invitrogen, Waltham, MA, USA) to enrich endothelial cells and eliminate contaminating fibroblasts. DMEC were routinely cultured in EGM-MV at 37°C, 5% CO_2_ and used for experiments between passage 3 and 8.

Patient-derived melanoma cell lines (Puujalka *et al*, 2016; Pirker *et al*, 2003), including VM15 (NRAS mutant), VM53 (BRAF and NRAS wildtype), VM21 and VM48 (both BRAF mutant) were cultured in RPMI-1640 (21875-034) supplemented with 10% FCS and 50 U/ml streptomycin-penicillin (15070-063), which were all from Gibco. All cells were maintained in a humidified atmosphere containing 5% CO_2_ at 37°C and passaged at 90% confluence.

### Western Blot

After treatment with the indicated BRAFi for 1 h, DMEC or melanoma cells were washed with ice-cold PBS and lysed on ice using a radioimmunoprecipitation buffer (RIPA, containing 50 mM Tris-HCl, 150 mM NaCl, 1% NP-40, 0.5% sodium deoxycholate, 0.1% SDS, and 1 mM EDTA) supplemented with protease inhibitor (P8340, Sigma-Aldrich) and phosphatase inhibitor cocktails (P5726, P0044, Sigma-Aldrich). Lysates were centrifuged at 18,000 g for 15 min at 4°C and supernatants were used for further analysis. Protein concentrations were determined using Bradford Protein assay (500-0006, Bio-Rad, Hercules, CA, USA) according to the manufacturer’s protocol. Per sample, 20 µg of protein was mixed with reducing Laemmli buffer and denatured for 5 min at 95°C. After SDS-PAGE, proteins were transferred onto a nitrocellulose membrane (GE Healthcare Life Sciences, Chicago, IL, USA) by wet blotting. Equal protein loading was confirmed by staining with Ponceau-S (33427, Serva, Heidelberg, Germany). After blocking with 5% bovine serum albumin (BSA, A2153, Sigma-Aldrich) in Tris-buffered saline buffer containing 0.1% Tween-20 (TBST), membranes were washed with TBST and incubated overnight at 4°C with the indicated primary antibodies diluted in TBST and 5% BSA. After washing with TBST, the membranes were incubated with fluorescent secondary antibodies (LI-COR), diluted in 5% milk powder (70166, Sigma-Aldrich) in TBST for 1 h at room temperature (RT). Blots were imaged and analyzed using Odyssey CLx and Image Studio (version 5.2) from LI-COR.

### Phosphoproteomics

#### Sample preparation and TMT labeling

Confluent DMEC from three donors were treated with the indicated inhibitor concentrations or vehicle control for 1 h. After washing with ice-cold PBS, cells were lysed on ice with 1% SDS and protease/phosphatase inhibitor cocktails. Lysates were homogenized by sonication. Lysates were incubated with Dithiothreitol (DTT) for 30 min at 37°C and alkylated with 20 mM iodoacetamide for another 30 min. Proteins were purified by precipitation with ethanol-acetone, and protein pellets were resolubilized in 1% sodium deoxycholate (SDC) in 50 mM Triethylammonium bicarbonate (TEAB) buffer, pH 8. For each sample, 150 µg proteins were brought to final volume of 95 µl (in SDC-TEAB buffer) and digested, firstly with Lysyl-endopeptidase (0.01 AU, FUJIFILM Wako Pure Chemicals Corp., Osaka, Japan) for 2 h at 37°C and subsequently with trypsin (7.5 µg per sample) for 4 h at 37°C. Peptides in each sample were labeled with one of the tags of TMTpro 18-plex labeling kit (A52045, Thermo Fisher Scientific) according to the manufacturer’s instructions. All 18 samples were then pooled in a 1:1 total TMT channel intensity ratio, measured by high resolution LC-MS2. Pooled samples were acidified (pH < 2) with formic acid to precipitate SDC and the collected supernatant was lyophilized.

#### TiO_2_ phosphopeptide enrichment

Phosphopeptides were isolated using TiO_2_ affinity chromatography as previously described (Engholm-Keller & Larsen, 2016). Briefly, dried peptides were dissolved in TiO_2_ loading buffer (80% acetonitrile (ACN), 5% Trifluoroacetic acid (TFA) and 1 M glycolic acid, all from Sigma) and incubated with 0.6 mg TiO_2_ beads (Titansphere, GL Sciences, Torrance, CA, USA) per 100 µg peptide solution for 30 min at RT on a shaker. The beads were spun down and supernatant was transferred to a new tube with 0.3 mg beads per 100 µg peptide. After 15 min shaking at RT, beads were spun down again, and supernatant was collected in a separate tube. TiO_2_ beads from both incubations were subsequently washed with 80% ACN/1% TFA and 10% ACN/0.1% TFA. The unbound peptides in the supernatants from incubation and wash steps were combined and stored as “unmodified peptides” (further details in supplementary methods). The TiO_2_ beads with bound phosphopeptides were resuspended in 100 µl of 100 mM TEAB, pH 8.5 and incubated with PNGase F (1,000 U, New England BioLabs, Ipswich, MA, USA) and Sialidase A (5 mU, Prozyme/Agilent, Santa Clara, CA, USA) overnight at 37°C in order to deglycosylate peptides (Larsen *et al*, 2007). The phosphorylated peptides were eluted from the TiO_2_ beads by incubation with 1.5% ammonium hydroxide solution, pH 11.3, for 10 min at RT with vigorous shaking. The beads were spun down and the supernatant was passed through a C8 membrane (3 M Empore™, Sigma-Aldrich), to remove any residual beads. The membrane was then washed with 100 µl of 50% ACN to obtain any retained peptide, before the samples were dried. To reduce sample complexity, high-pH reversed-phase fractionation was applied (Boll *et al*, 2020). Phosphopeptides were dissolved in 20 mM ammonium formate, pH 9.3, and loaded on an Acquity UPLC^®^-Class CSHTM C18 column (Waters, Milford, MA, USA). Fractionation was performed on a Dionex Ultimate 3000 HPLC system (Thermo Fisher Scientific), and a total number of 20 concatenated fractions was collected.

#### Nano-flow liquid chromatography-mass spectrometry (nLC-MS/MS) analysis

The analysis was performed on an Easy-nLC System (Thermo Fisher Scientific) using buffer A (0.1% formic acid, FA) and buffer B (95% ACN, 0.1% FA) and an Orbitrap Eclipse Tribrid MS (Thermo Fisher Scientific). All fractions were redissolved in buffer A (0.1% FA) and loaded into the in-house made fused silica capillary column setup (18 cm pulled emitter analytical column with 75 μm inner diameter, packed with Reprosil-Pur 120 C18-AQ, 3 μm (Dr. Maisch GmbH, Ammerbuch, Germany)). The peptides were eluted with gradient elution from 2% to 95% buffer B, with a flow rate of 300 nl/min. Ionization was performed by nano-electrospray. Phosphopeptides were analyzed by data-dependent acquisition in positive ion mode mass spectrometry. The m/z scan range for full MS scan was 350 – 1500 Da, and intact peptides were detected in the Orbitrap with a resolution of 120,000 full width half maximum (FWHM), a normalized Automatic gain control (AGC) target value of 250%, and a maximum injection time of 50 ms. From each full scan, the top 10 most intense precursor ions were selected for higher energy collision dissociation (HCD) fragmentation with a normalized collision energy (NCE) of 35%. The MS^2^ was performed with the following parameters: orbitrap resolution of 45,000 FWHM, normalized AGC target value of 300%, isolation window of 1.2 m/z, dynamic exclusion window of 3 s and a maximum injection time in automatic mode.

#### Protein identification and quantification

Protein identification was performed using Proteome Discoverer (version 2.4.0.305, Thermo Fisher Scientific). The search was performed against the UniProtKB/Swissprot database (homo sapiens, release-2022_04/) using an in-house Mascot server (v2.8.2, Matrix Science Ltd, London, UK) and the built-in Sequest HT search engine. Fixed modifications in the search included TMT-Pro_K (K), TMT-Pro_N term (N-term) and Carbamidomethyl (C), whereas Deamidated (N) and Phosphorylation (S, T, Y) were set as dynamic modifications. Further parameters of the search included a fragment mass tolerance of 0.03 Da, a precursor mass tolerance of 10 ppm and maximum of two missed cleavages. Data filtering was performed using a percolator, with ≤ 1% false discovery rate (FDR) (peptide and protein level). This resulted in a list of 7,756 master proteins, 11,458 peptide groups (filtered for phosphorylation modification), 12,543 peptide isoforms, 187,269 PSMs and 1,022,760 MS/MS spectra. The abundance values of peptide groups were normalized to the total peptide amount of each channel through Proteome Discoverer. Since the treatments did not cause changes in cellular protein expression but only resulted in minor losses of extracellular matrix components in the 100 µM Vemurafenib samples (see Figure S2) phosphopeptide abundances were not further adjusted to respective protein levels.

#### Bioinformatic analysis

Differences between treatments and the vehicle control were determined via Limma testing (including paired tests within donors), using the combined statistical testing tool PolySTest (Schwämmle *et al*, 2020). Phosphosites were considered as significantly altered at a log^2^-fold change ± 1 and an FDR ≤ 0.05. Reactome (v84, reactome.org) pathway enrichment analysis of significantly altered proteins was performed for each treatment. The resulting lists of pathways were then filtered for hits where at least one treatment fulfilled the thresholds (p ≤ 0.05 and strength [entities found / entities in the pathway] ≥ 0.05). Terms involving infectious disease (R-HSA-1169410, R-HSA-9609690, R-HSA-8875360, R-HSA-1169408, R-HSA-8876384) were removed due to contextual inapplicability. Data visualization was performed with R version 4.2.2 (R Foundation for Statistical Computing, Vienna, Austria). Additionally, kinase activity prediction was performed using the KinSwingR package, based on phosphopeptide abundance in our dataset and known kinase-substrate interactions from the PhosphoSitePlus database (Engholm-Keller *et al*, 2019). Physical subnetwork visualizations of the 50 most differentially regulated kinases among all treatments were created with the Cytoscape software (version 3.9.1), including the applications Omics Visualizer and STRING, using a 0.6 confidence score cutoff.

### Electrical cell-substrate impedance sensing (ECIS)

ECIS (Applied Biophysics, Troy, NY, USA) was used to measure barrier resistance of DMEC monolayers. 8W10E+ array plates (72040, ibidi, Planegg, Germany) were coated with 1% gelatin before cell seeding at a density of 15,000 DMEC/cm². Resistance was measured continuously in a multi-frequency setup. After the resistance at 4,000 Hz reached a stable plateau of > 1,000 Ω, endothelial cells were treated and continuously monitored at 250 Hz as previously described (Schossleitner *et al*, 2016).

### Permeability of fluorescent tracers

DMEC were seeded into transwell inserts (734-2747, VWR International, Radnor, PA, USA) and cultured until confluence. Indicated treatments were added to the transwell, along with 0.2 µg/ml Na-Fluorescein (376 Da, F6377, Sigma-Aldrich) and 50 µg/ml TRITC-conjugated dextran (70 kDa, D1818, invitrogen) tracers. At indicated timepoints, fluorescence intensity was measured in the medium below the transwell insert with a standard plate reader, using the settings for Fluorescein (excitation: 485 nm, emission: 535 nm) and TRITC (excitation: 540 nm, emission: 600 nm).

### Immunofluorescence

DMEC were seeded on µ-Slide chamber slides (ibidi), grown to 100% confluence, and fixed with 4% paraformaldehyde (PFA) for 15 min at RT. Following permeabilization in 100% methanol for 10 min at −20°C, cells were stained with indicated primary antibodies diluted in PBS containing 1% BSA overnight at 4°C and appropriate secondary antibodies for 1 h at RT. Nuclei were stained with DAPI (D9542, 1:1000, Sigma-Aldrich). Cells were then imaged using a confocal laser scanning microscope (LSM-980; Carl Zeiss, Jena, Germany) equipped with a Plan-Apochromat 63×/1.40 oil lens.

### Melanoma spheroid-induced gap formation

Based on a previously published in vitro assay of tumor cell invasion (Holzner *et al*, 2016), BRAF-mutant VM48 melanoma cells (1,500 per well) were seeded in a round-bottom 96-well plate (Greiner Bio-One, Monroe, NC) in full RPMI containing 0.3% methylcellulose (4,000 cP; M0512, Sigma-Aldrich), followed by centrifugation for 15 min at 1,200 rpm and 12°C, and incubation for 3-4 days. Meanwhile, DMEC were seeded on µ-Slide 4-well chamber slides (ibidi) and grown to 100% confluence. After staining with CellTracker green CMFDA Dye (C2925, Invitrogen), DMEC were pre-treated with the indicated inhibitor concentrations for 6 h before washing with medium. Subsequently, melanoma spheroids were collected, carefully washed and resuspended in EGM2-MV and added onto the endothelial monolayer (approx. 24 spheroids per well). After an incubation of 6 h, chamber slides were scanned with an automated microscope (Cytation 5, Agilent) with a 4x objective and filters for high-contrast brightfield and GFP fluorescence. The area of circular discontinuities within the endothelial monolayer beneath the spheroids was quantified using FIJI software (Fiji is just ImageJ, version 1.54b).

### Histology and immunofluorescence of patient samples

We conducted a comprehensive screening of archived histological samples from melanoma patients who visited the Department of Dermatology at the Vienna General Hospital between 2012 and 2022. The study was conducted according to the principles expressed in the Declaration of Helsinki. We identified a total of 90 patients who met our predefined criteria, as outlined in the ethical protocol approved by the ethics committee of the Medical University of Vienna with approval number 1820/2022. Inclusion criteria were defined as follows: i) verified diagnosis of melanoma stage IIIA-IVM1d, ii) BRAF-V600E/K mutation, iii) therapy with either Vemurafenib alone, Vemurafenib + Cobimentinib, or Dabrafenib + Trametinib, and iv) an age of 18-99 years at the time of sample collection. Patients were excluded from the study if they received simultaneous treatment with other cancer therapies such as ICIs. Upon further evaluation, only 15 patients had available matching pre- and on-treatment cutaneous metastatic tissue samples. Out of those, we excluded nine patients who did not have skin biopsies available, but metastatic tissue samples from other organs. One sample was not released for research purposes. Consequently, we included a total of five patients, four of which were unique individuals and one was a matched pair based on age, sex, disease stage, and lactate dehydrogenase (LDH) levels. The patient cohort included three males and two females, with a median age of 70 years (range: 39 – 81 years) and a median LDH level of 208 U/l (range: 183 – 484 U/l). Their disease stages were classified as IVM1c (n = 4) or IVM1d (n = 1) according to the American Joint Committee on Cancer (AJCC) 8^th^ edition staging system. All included melanomas were tested positive for the BRAF-V600E mutation and had been treated either with Vemurafenib monotherapy (n = 1), Vemurafenib + Cobimetinib (n = 1), or Dabrafenib + Trametinib (n = 3). From the patients’ formalin-fixed paraffin-embedded (FFPE) cutaneous metastatic tissue samples, sections with a thickness of 7 µm were cut and stained via immunofluorescence for the indicated antibodies (see Table 3), nuclei were counterstained with DAPI. Slides were scanned using a Vectra Polaris imaging system (Akoya Biosciences, Inc., Marlborough, MA, USA) with a 20x objective. Image analysis and fluorescence intensity quantification was performed in peritumoral tissue areas via the QuPath software (version 0.4.3) (Bankhead *et al*, 2017).

### Statistical rationale

Unless otherwise specified, differences between treatments and the control were analyzed using one-way ANOVA, corrected with Dunnett’s multiple comparisons test in GraphPad Prism (version 8.0.1). Significance levels are depicted in the graphs as follows: p < 0.05 (*), p < 0.01 (**), p < 0.001 (***), p < 0.0001 (****).

## Data availability

The mass spectrometry data and Proteome Discoverer files from this publication will be deposited to the PRIDE database [www.ebi.ac.uk/pride/] and assigned the identifier [accession]. Imaging data will be available at Imaging Data Resource [idr.openmicroscopy.org] under the identifier [accession].

## Acknowledgements

This work has been funded by the Vienna Science and Technology Fund (WWTF) [10.47379/LS18080] to KS, by the Comprehensive Cancer Center Vienna (Cancer Research Initiative Grant to SB), the European Molecular Biology Organization (Scientific Exchange Grant #9313 to SB), the City of Vienna (Cancer Research Fund #22077, to KS and SB). Images were obtained at the Core Facility Imaging of the Medical University Vienna. We thank Marion Gröger (†), Sabine Rauscher, Christoph Friedl and Philipp Velicky for their continuous support. This study was supported by the Villum Center for Bioanalytical Sciences at University of Southern Denmark.

## Author contributions

**Sophie Bromberger:** Conceptualization; data curation; formal analysis; funding acquisition; investigation; methodology; project administration; visualization; writing (original draft); writing (review and editing). **Yuliia Zadorozhna:** Formal analysis; investigation; writing (review and editing). **Julia Maria Ressler:** Methodology; resources; writing (review and editing). **Silvio Holzner:** Methodology. **Arkadiusz Nawrocki:** Methodology; writing (review and editing). **Nina Zila:** Methodology; writing (review and editing). **Alexander Springer:** Resources. **Martin Røssel Larsen:** Methodology; resources; supervision; writing (review and editing). **Klaudia Schossleitner:** Conceptualization; funding acquisition; methodology; project administration; supervision; writing (original draft); writing (review and editing).

