## Supplement for "Off-targets of BRAF inhibitors disrupt endothelial signaling and differentially affect vascular barrier function"

### Supplementary methods

##### Assessment of endothelial morphology

DMEC were seeded on µ-Slide chamber slides (ibidi) and grown to 100% confluence, before staining with CellTracker green CMFDA Dye (C2925, Invitrogen) according to the manufacturer's instructions. Cells were treated with the indicated inhibitor concentrations and scanned after 0, 1 and 6 h with an automated microscope (Cytation 5, Agilent) with a 10x objective and filters for phase contrast and GFP fluorescence.

##### Endothelial adhesion markers

DMEC were seeded onto 24-well plates and grown to confluence, before being treated with the indicated inhibitor concentrations or 100 ng/ml bacterial lipopolysaccharide (LPS, L2280, Sigma-Aldrich) for 6 h. After washing cells with PBS, they were detached with trypsin. Cell suspensions were washed with PBS and with 1% BSA. Each suspension was stained with directly labelled antibodies for ICAM-1 (PE-conjugated, 1:100, 347970, BD Biosciences, Franklin Lakes, NJ, USA) and E-Selectin (FITC-conjugated, 1:50, 61R-CD62ebHUFT, Fitzgerald, Acton, MA, USA) for 1 h at 4°C. Suspensions were washed with PBS before measurements on a CytoFLEX flow cytometer (Beckman Coulter). Data were analyzed in the CytExpert software (version 2.5).

##### Proteomics of unmodified peptides

After enrichment of phosphopeptides, supernatants from incubation and wash steps were collected and stored as the unmodified fraction. The unmodified peptides were purified using a commercial HLB cartridge that was equilibrated with ACN and 0.1% TFA, subsequently, before loading and washing the sample with 0.1% TFA. Elution was performed with 65% ACN, 0.1% TFA solution, and samples were dried. Simultaneously to the phosphopeptides, unmodified peptides were also dissolved in buffer A and loaded on an Acquity UPLC®-Class CSHTM C18 column (Waters, USA). The fractionation was performed on a Dionex Ultimate 3000 HPLC system (Thermo Scientific, USA), and 20 concatenated fractions were collected.

Analysis of the unmodified peptides was performed with the same instrumentation as the phosphopeptides. The peptides were eluted with gradient elution using a flowrate of 250 nl/min from 2% to 95% buffer B and were analyzed by data-dependent acquisition and positive ion mode mass spectrometry. The m/z scan range for full MS scan was 350 – 1600 Da, and intact peptides were detected in the Orbitrap with a resolution of 120,000 FWHM, an AGC target value of 1 × $10^6$ ions, and a maximum injection time of 50 ms. Further, the peptides were selected for collision-induced dissociation (CID), using fixed collision energy set to 35%. The fragment ion spectra were recorded using the Ion trap at a resolution of 15,000, an AGC target value of 1 × $10^5$ $MS^2$ ions and maximum injection time of 150 ms. $MS^3$ spectra were acquired using synchronous precursor selection (SPS) of 10 precursors. $MS^3$ precursors were fragmented by HCD with 55% of the collision energy and analyzed using the Orbitrap at 30,000 resolution power with a scan range of 100 - 500 m/z, an isolation window of 2 m/z, an AGC target value of 1 × $10^5$ $MS^2$, and maximum injection time of 120 ms with one microscan.

Protein identification and quantification was performed using Proteome Discoverer, as described above. The search resulted in 8,132 master proteins, 49,200 peptide groups, 73,834 peptide isoforms, 176,894 PSMs and 1,523,663 MS/MS spectra. For statistical testing the combined statistical testing tool PolySTest was used and data were visualized with R. Normalized abundances of proteins were analyzed via Limma testing including paired tests within donors. Differences between treatments and the vehicle control were considered statistically significant with $log^2$-fold change ± 1 and an FDR ≤ 0.05.

### Supplementary Figure legends

**Figure S1: BRAFi differentially affect ERK activation in DMEC and melanoma cells.**

A: Abundances of pERK1/2 (T202/Y204), total ERK1/2 and GAPDH in DMEC and melanoma cells after 1 h of indicated BRAFi treatment.

**Figure S2: BRAFi do not affect overall protein expression.**

Mass spectrometry-based proteomics data of DMEC treated with the indicated inhibitors for 1 h (same samples as in Figure 2).

A-E: Protein abundances of treated cells compared to vehicle control (Limma, logFC ± 1, $p \leq 0.05$).

F: Quantification of differentially recovered peptides compared to vehicle control.

**Figure S3: BRAFi do not affect endothelial morphology nor the surface expression of activation markers.**

A: Fluorescence images of DMEC, stained with green CellTracker, after 0, 1, and 6 h of BRAFi treatment.

B: Gating for endothelial activation markers ICAM-1 and E-Selectin, using unstained cells, or stained cells treated with 0.1% DMSO, or 10 ng/ml LPS as an example.

C: Percentage of ICAM-1 or E-Selectin positive cells upon treatment with 0.1% DMSO, 10 ng/ml LPS, or 1-100 µM BRAFi for 6 h.

**Figure S4: BRAFi differentially affect endothelial tight and adherens junctions**

Immunofluorescence images of confluent DMEC treated with the indicated inhibitors for 1 h. Green = Claudin-5, red = VE-Cadherin. Scale bars = 20 µm.

### Figure S1

A

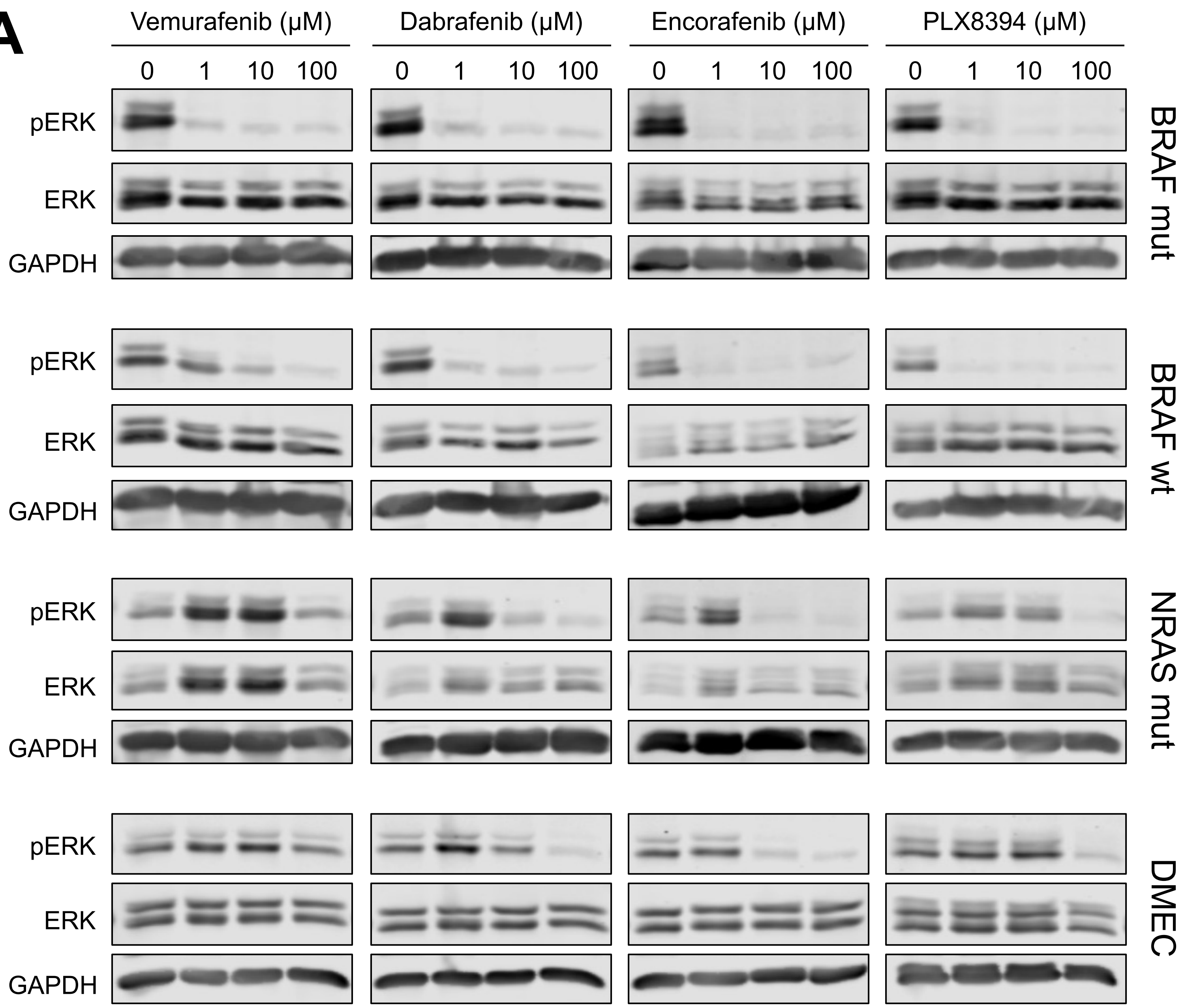

Vemurafenib (μM)
Dabrafenib (μM)
Encorafenib (μM)
PLX8394 (μM)
0 1 10 100
pERK
ERK
GAPDH
BRAF mut
BRAF wt
NRAS mut
DMEC

### Figure S2

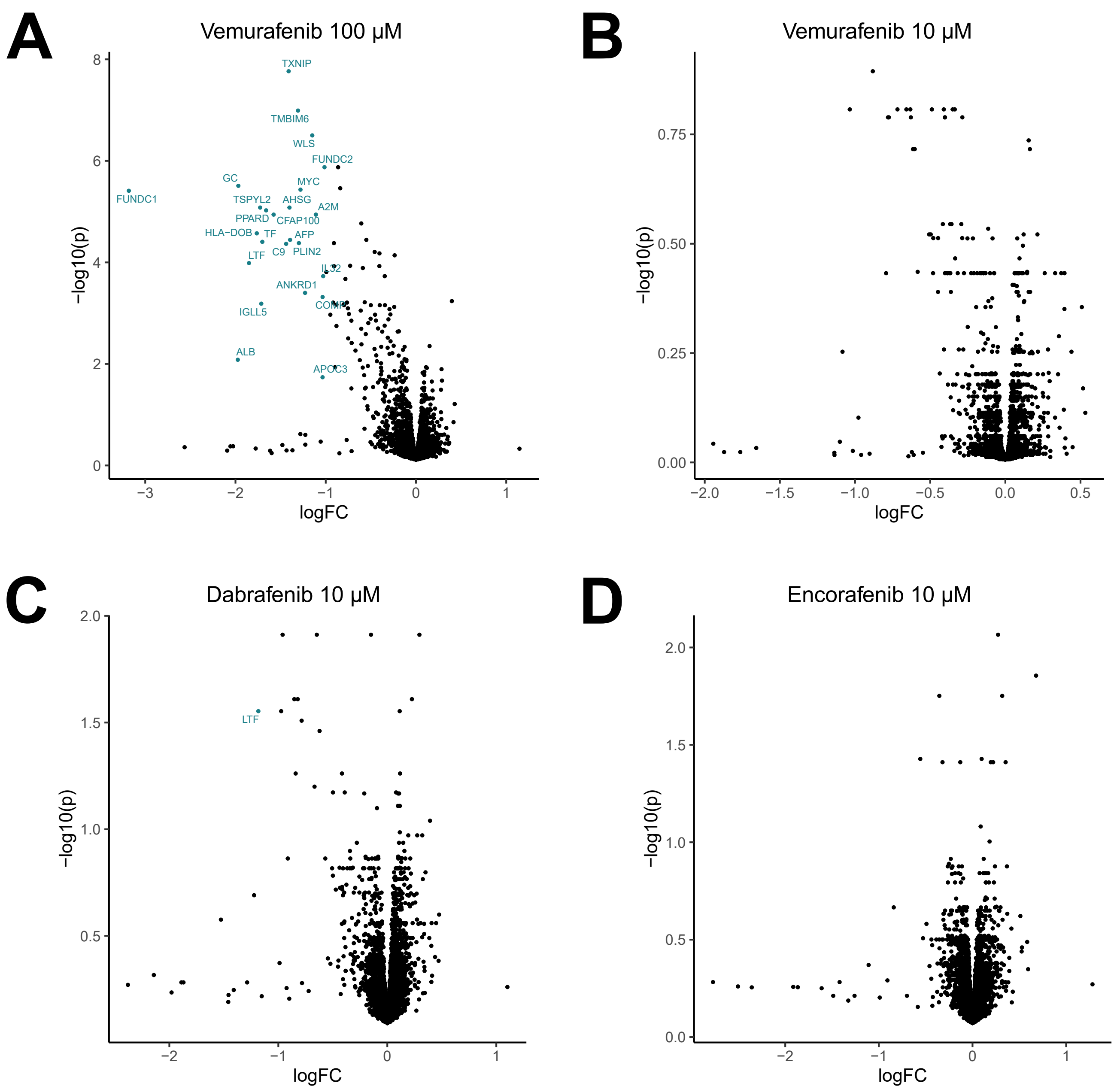


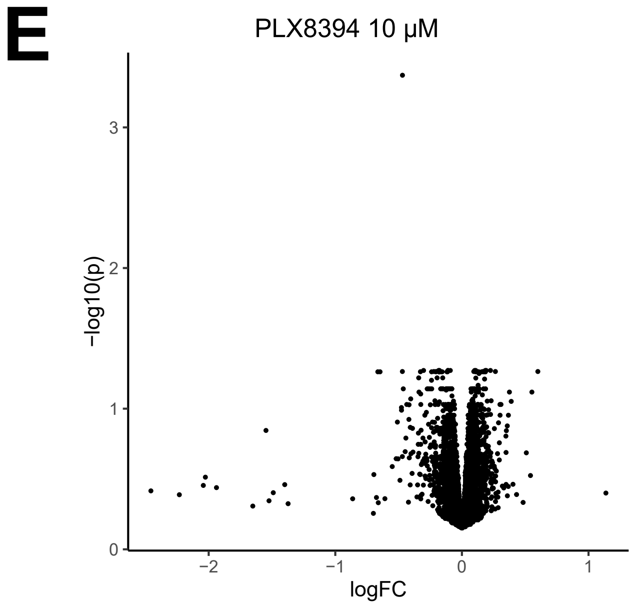


**F**

| proteins (p ≤ 0.05) | | | |
|---|---|---|---|
| **Treatment** | **logFC < -1** | **logFC > 1** | **total** |
| V100 | 24 | 0 | 24 |
| V10 | 0 | 0 | 0 |
| D10 | 1 | 0 | 1 |
| E10 | 0 | 0 | 0 |
| P10 | 0 | 0 | 0 |

### Figure S3

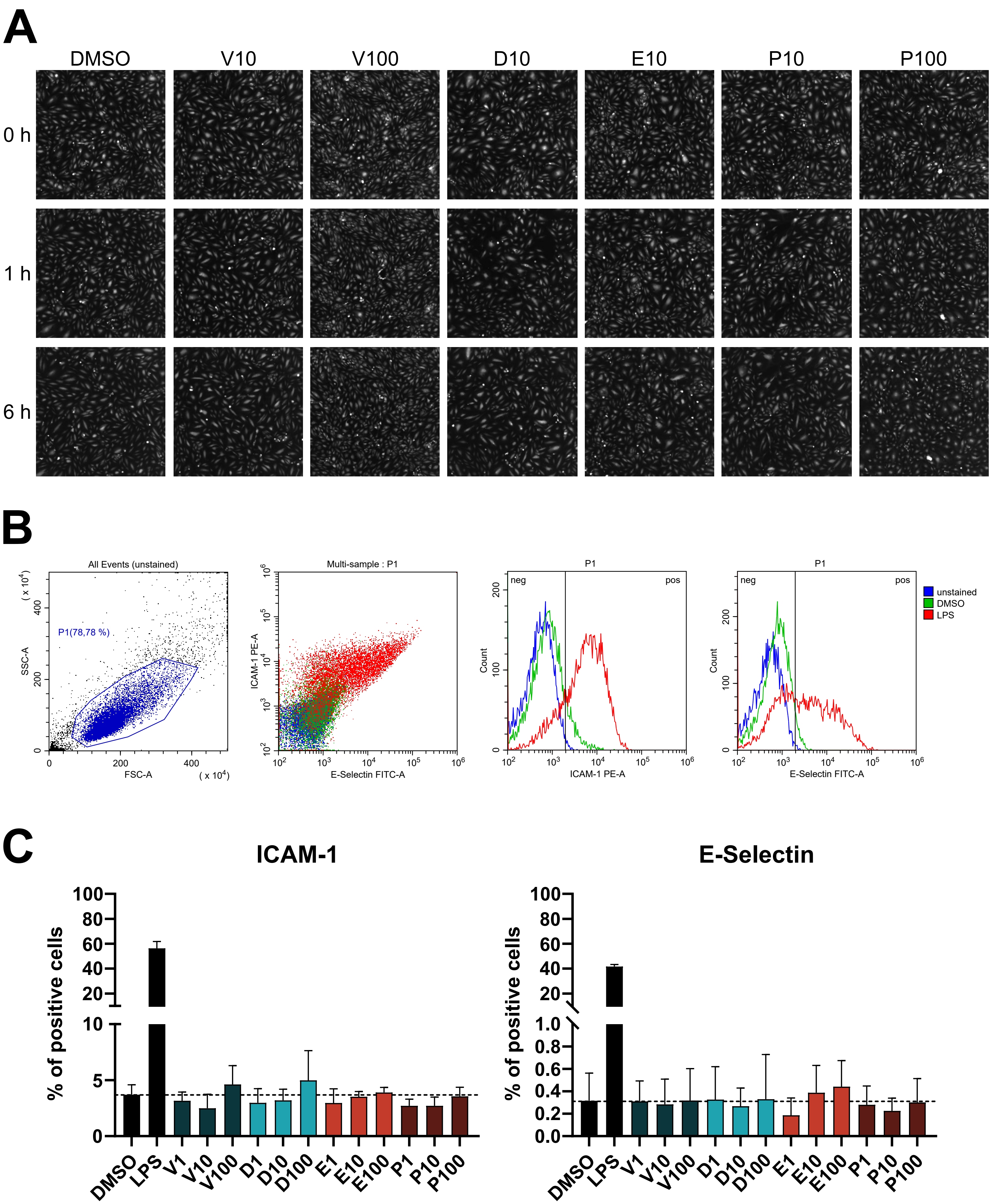
A
DMSO
V10
V100
D10
E10
P10
P100
0 h
1 h
6 h
B
All Events (unstained)
P1(78,78 %)
SSC-A
FSC-A
( x 10^4)
Multi-sample : P1
ICAM-1 PE-A
E-Selectin FITC-A
P1
neg
pos
Count
ICAM-1 PE-A
E-Selectin FITC-A
unstained
DMSO
LPS
C
ICAM-1
E-Selectin
% of positive cells
DMSO
LPS
V1
V10
V100
D1
D10
D100
E1
E10
E100
P1
P10
P100

### Figure S4

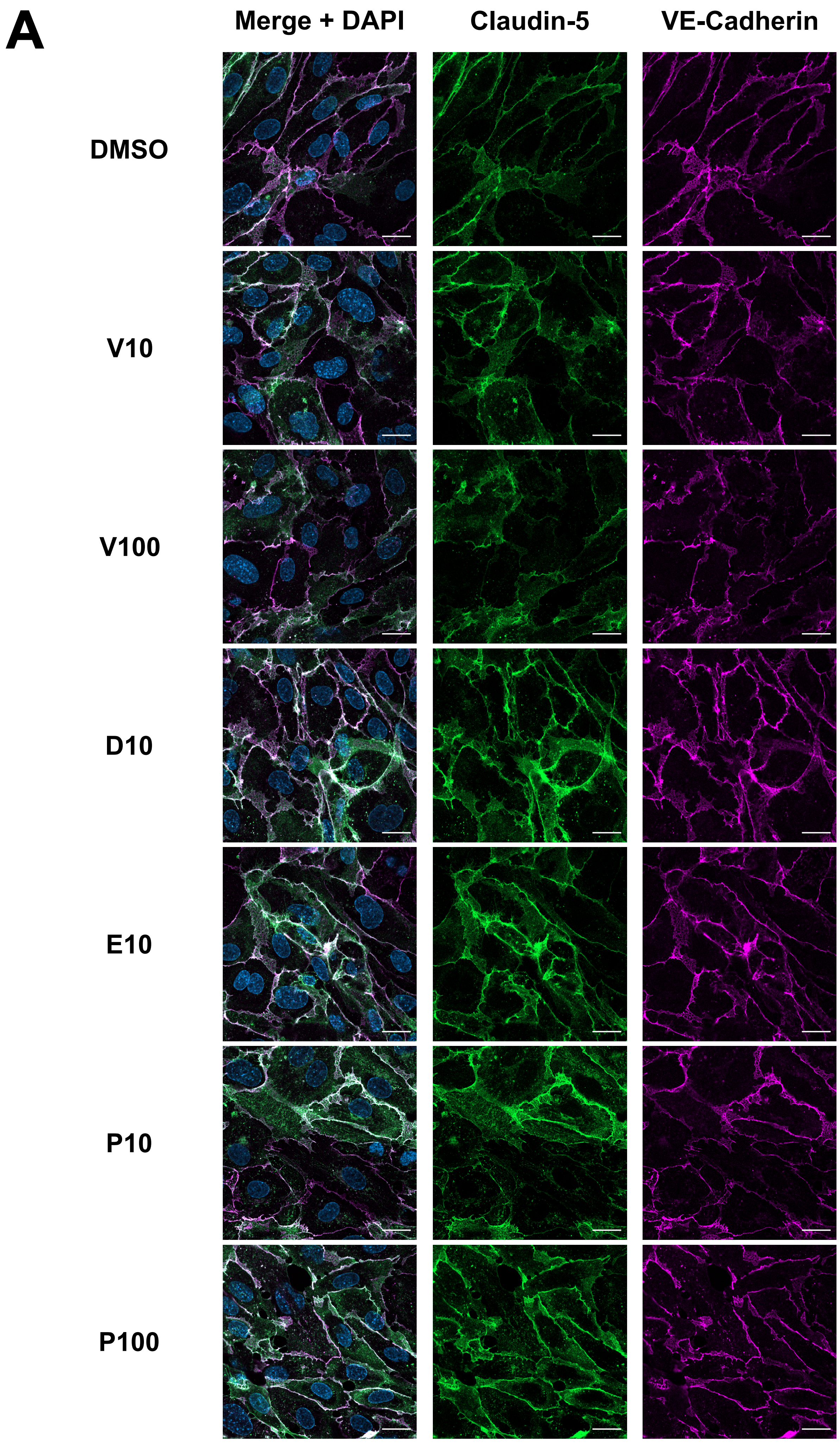
A
Merge + DAPI
Claudin-5
VE-Cadherin
DMSO
V10
V100
D10
E10
P10
P100
